# Actin fence therapy with exogenous V12Rac1 protects against Acute Lung Injury

**DOI:** 10.1101/2021.01.20.427469

**Authors:** Galina A. Gusarova, Shonit R. Das, Mohammad N. Islam, Kristin Westphalen, Guangchun Jin, Igor O.Shmarakov, Li Li, Sunita Bhattacharya, Jahar Bhattacharya

**Author notes:** To whom correspondence should be addressed: Jahar Bhattacharya, M.D., D.Phil., Department of Medicine, 630 W. 168th Street, Room BB 8-812, New York, NY 10032. Deceased 4 Nov 2018 (Shonit R. Das).

## Abstract

High mortality in Acute Lung Injury (ALI) results from sustained proinflammatory signaling by alveolar receptors, such as TNFα receptor type 1 (TNFR1). Factors that determine the sustained signaling are not known. Unexpectedly, optical imaging of live alveoli revealed a major TNFα-induced surge of alveolar TNFR1 due to a Ca^2+^-dependent mechanism that decreased the cortical actin fence. Mouse mortality due to inhaled LPS was associated with cofilin activation, actin loss and the TNFR1 surge. The constitutively active form of the GTPase, Rac1 (V12Rac1), given intranasally as a non-covalent construct with a cell-permeable peptide, enhanced alveolar F-actin and blocked the TNFR1 surge. V12Rac1 also protected against ALI-induced mortality resulting from intranasal (i.n.) instillation of LPS, or of *Pseudomonas aeruginosa*. We propose a new therapeutic paradigm in which actin enhancement by exogenous Rac1 strengthens the alveolar actin fence, protecting against proinflammatory receptor hyperexpression, hence blocking ALI.

## Introduction

Life-threatening tissue injury to critical organs occurs as a result of host-pathogen interactions involving proinflammatory receptors. In lung, the resulting inflammation underlies acute lung injury (ALI), which can lead to the acute respiratory distress syndrome (ARDS), a condition that associates with high mortality (1). Pharmacological therapies are not available for ALI, but are required in order to stem disease progression.

ALI due to inhaled Gram-negative bacteria occurs through initiating and progressive phases. In the initiating phase, inhaled pathogens ligate toll-like receptors on macrophages and the alveolar epithelium (2, 3) causing release of proinflammatory cytokines, such as TNFα that ligate alveolar epithelial receptors (4). Crosstalk with the endothelium follows (5) and chemoattractants, such as IL-8 activate inflammatory cell recruitment (6). In the progressive phase, recruited and resident immune cells continue to secrete cytokines thereby sustaining the inflammatory response (7, 8). The extent to which the initiating mechanisms continue to enhance the progressive phase of the response remains unclear. Here we addressed this question in the context of the interactions of TNFα with its alveolar receptor, TNFR1.

Studies from multiple cell types indicate that following ligation, TNFR1, a transmembrane protein, sheds its ectodomains (9–11). The sheddase, ADAM-17 mediates the shedding (10), which is protective since shedding inhibition augments lung inflammation (9). The shed domains are not recycled by the cell from which the shedding occurred (12). Brefeldin A, the inhibitor of protein trafficking to the Golgi, abrogated TNFR1 receptor mobilization and decreased TNFR1 abundance on the cell surface, indicating that membrane replenishment of TNFR1 occurs by receptor trafficking from storage pools in the Golgi (11, 13–15). The replenishment is translation independent (13, 16). However, the extent and the time course of TNFR1 replenishment in the alveolar epithelium remain unknown.

In this regard, the regulatory role of cortical actin remains unclear. Cortical actin is the layer of filamentous actin (F-actin) that forms a network adjacent to the plasma membrane (PM). Although cortical actin is known to form a “fence” against trafficking of vesicles and receptors to the PM (17, 18), dynamic data from live alveoli are lacking detailing the role of the actin fence in TNFR1 expression. Here, we addressed these issues by application of confocal microscopy of the live alveolar epithelium. Our goals were to determine strategies by which fence enhancement might impede alveolar receptor display, hence ALI pathogenesis. We show for the first time, to our knowledge, that loss of the fence causes a proinflammatory surge of TNFR1 expression, and that the fence is a druggable target for mitigating ALI.

## Results

### F-actin determines alveolar TNFR1 display

Optical access to the live alveolar epithelium of mouse lung provided an opportunity for dynamic evaluation of the role of the F-actin fence as a determinant of epithelial TNFR1 expression. By real-time confocal microscopy of pulmonary alveoli, we determined fluorescence expressions of TNFR1 in the alveolar epithelium in terms of a fluorescent mAb that detects TNFR1 ectodomains (9) (Figure 1A). We confirmed that the TNFR1 immunofluorescence was on the cell surface, since it was eliminated by alveolar injection of a cell-impermeable fluorescence quenching agent. Our procedures did not label TNFR1 on the capillary endothelium adjacent to the epithelium (Supplemental Figure 1A). Moreover, co-staining with the lamellar body marker, Lysotracker Red (LTR) revealed TNFR1 expression on the type 1 epithelial cells (AT1), and no detectable expression of TNFR1 on the type 2 (AT2) (Supplemental Figure 1B).

**Figure 1.**
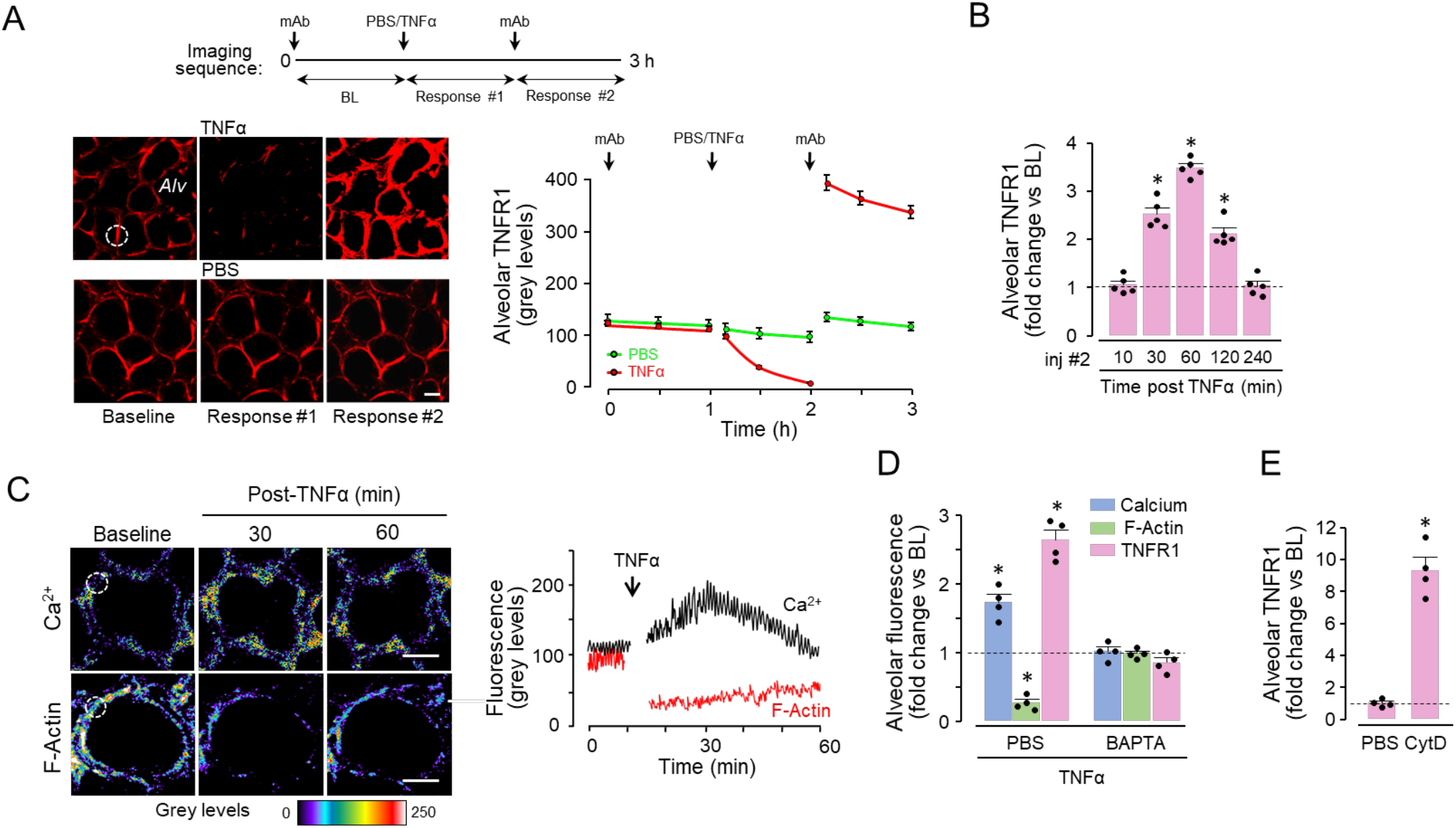
F-actin determines surge in alveolar TNFR1 expression. (**A**) Confocal images of live alveoli from single experiments show red immunofluorescence (IF) of TNFR1. The sketch summarizes the imaging protocol over a 3-hour period. After alveolar microinfusion of mAb (*1^st^ arrow*), baseline (*BL*) images were obtained every 30 minutes for 1 hour. Then, TNFα or PBS was given by alveolar microinfusion (*2^nd^ arrow*), and the response imaged at a similar rate (*Response #1*). Finally, the mAb was microinjected in the same alveoli for a second time (*3^rd^ arrow*), and a third set of images were obtained (*Response #2*). Example images from the three imaging sequences are shown. Fluorescence was quantified throughout the imaging period at a region of interest (ROI) selected on a baseline image (*dashed circle*). These quantifications are shown as the timeline plot with the timepoints of the microinjections marked (*arrows*). The plot shows steady baseline fluorescence for 1 hour, ruling out photobleaching. The middle-upper image and the plot show that IF was almost undetectable 1 hour after TNFα. However, as shown in the right-upper image and the plot, in the same TNFα-treated alveoli that had previously undergone IF loss, the second mAb injection revealed marked IF increase. PBS caused no changes. Each plotted point shows the average for 15 ROIs from 5 alveoli of a single lung (mean±SE). Replicated in 3 lungs. Scale bar, 20 μm. *Alv*, alveolus. (**B**) Data show group responses to alveolar microinjection of TNFα. The response time points are as indicated. For each bar, *n*=5. \**p*<0.05 vs baseline (dotted line). (**C**) Confocal images of live alveoli show pseudocolored fluorescence of cytosolic Ca^2+^ and F-actin, as determined by fluorescence of Fluo4 and Lifeact, respectively. The tracings show alveolar epithelial responses to alveolar microinfusion of TNFα in a single lung. Replicated 5 times in each of 3 lungs. For the tracings, in the baseline image, ROIs were selected at grey levels of 80-120 to accommodate detection of subsequent increases or decreases of fluorescence within the dynamic range of the imaging system. Scale bars, 10 μm. (**D,E**) Data are whole-image grey levels normalized to the corresponding baseline *(dashed lines)*. Responses shown are to alveolar microinjection of TNFα after 30 minutes. The responses to cytochalasin D (*CytD*) were obtained after 60 minutes (**E**). Bars are mean±SE. Each dot shows data for a single lung. For each bar, *n*=4 (**C, E**), or 5 (**D**) lungs. \**p*<0.05 vs baseline (dotted lines) using ANOVA with Bonferroni correction.

To determine the dynamics of epithelial TNFR1 expression, in each alveolar field we sequentially microinjected the anti-TNFR1 mAb, TNFα (or PBS), and then repeated the anti-TNFR1 Ab for a second time (Figure 1A). The first mAb injection marked the baseline TNFR1 expression on the alveolar epithelium. Alveolar microinjection of TNFα rapidly induced a time dependent decrease of TNFR1 immunofluorescence (Figure 1A), affirming our previous findings that TNFα causes shedding of TNFR1 ectodomains (9). However, in the same alveoli that had undergone receptor shedding, the second mAb injection revealed marked enhancement of TNFR1 re-expression (Figure 1A).

In separate experiments, we gave the second mAb injection at different time points to further delineate the time course of TNFR1 re-expression. Our findings indicate that within 10 minutes after the TNFα-induced shedding, TNFR1 expression was similar to baseline (Figure 1B), indicating that the receptor was rapidly replaced. Subsequently, there was a surge of TNFR1 re-expression that on average, reached a peak at 1 hour and returned to baseline at 4 hours (Figure 1B).

To determine the effects of the cytosolic Ca^2+^ on F-actin and TNFR1 expression, we microinfused TNFα in alveoli expressing a transfected F-actin probe (5). At baseline, fluorescence of Ca^2+^ and F-actin were steady for at least 20 min (Figure 1C), ruling out photobleaching as an artefact. Alveolar microinjection of TNFα rapidly increased Ca^2+^, while concomitantly decreasing F-actin (Figure 1C). To inhibit the Ca^2+^ response, we gave alveolar microinjection of the Ca^2+^ chelator, BAPTA-AM, then determined responses after 30 minutes. BAPTA-AM blocked the TNFα-induced cytosolic Ca^2+^ increase, the F-actin decrease and the TNFR1 surge (Figure 1D). PBS pre-treatment had no effect. These first dynamic quantifications of actin in live alveoli indicated a strong effect of the cytosolic Ca^2+^ on F-actin and TNFR1 expression.

To depolymerize F-actin by a receptor-independent mechanism, we gave alveolar microinjection of the actin depolymerizing agent, cytochalasin D (cytD) and determined the TNFR1 response after 60 min (19). CytD exposure markedly enhanced TNFR1 expression (Figure 1E). Thus, even in the absence of TNFα, actin depolymerization with CytD was sufficient to induce the TNFR1 surge. Taken together, these findings affirmed that F-actin constitutively inhibited alveolar TNFR1 expression, and that decrease of F-actin caused the TNFR1 surge.

### TNFα causes calcineurin dependent alveolar TNFR1 expression

To determine mechanisms downstream of the TNFα-induced cytosolic Ca^2+^ increase, we considered the role of calcineurin (Cn), which is a Ca^2+^ sensor. Cn is a Ca^2+^-calmodulin dependent protein phosphatase that contains the catalytic CnA (α, β and γ isoforms) and the Ca^2+^-sensing CnB subunits (20, 21). In CnAβ-null mice, the β isoform of CnA is disrupted, inhibiting Cn’s phosphatase activity (22). Optical imaging revealed that the TNFα-induced F-actin decrease and the accompanying TNFR1 surge were absent in CnAβ-null mice (Figure 2A, Supplemental Figure 2A). Moreover, the calcineurin inhibitor, FK-506 blocked the TNFR1 surge (Supplemental Figure 2B). These findings mechanistically implicated calcineurin in the TNFR1 responses. Calcineurin dephosphorylates, hence activates the actin-severing protein, cofilin (23, 24). To evaluate this hypothesis, we carried out immunoblots on lysates of lungs of wild type and CnAβ-null mice. Our findings indicated that levels of phosphorylated (inactive) cofilin were higher in CnAβ-null than WT mice (Figure 2B). Moreover, 4 hours after intranasal (i.n.) instillation of the TNFα, cofilin dephosphorylation occurred in wild type (WT), but not CnAβ-null mice (Figure 2B). We interpret from these findings that TNFα-induced calcineurin activation, led to cofilin dephosphorylation.

**Figure 2.**
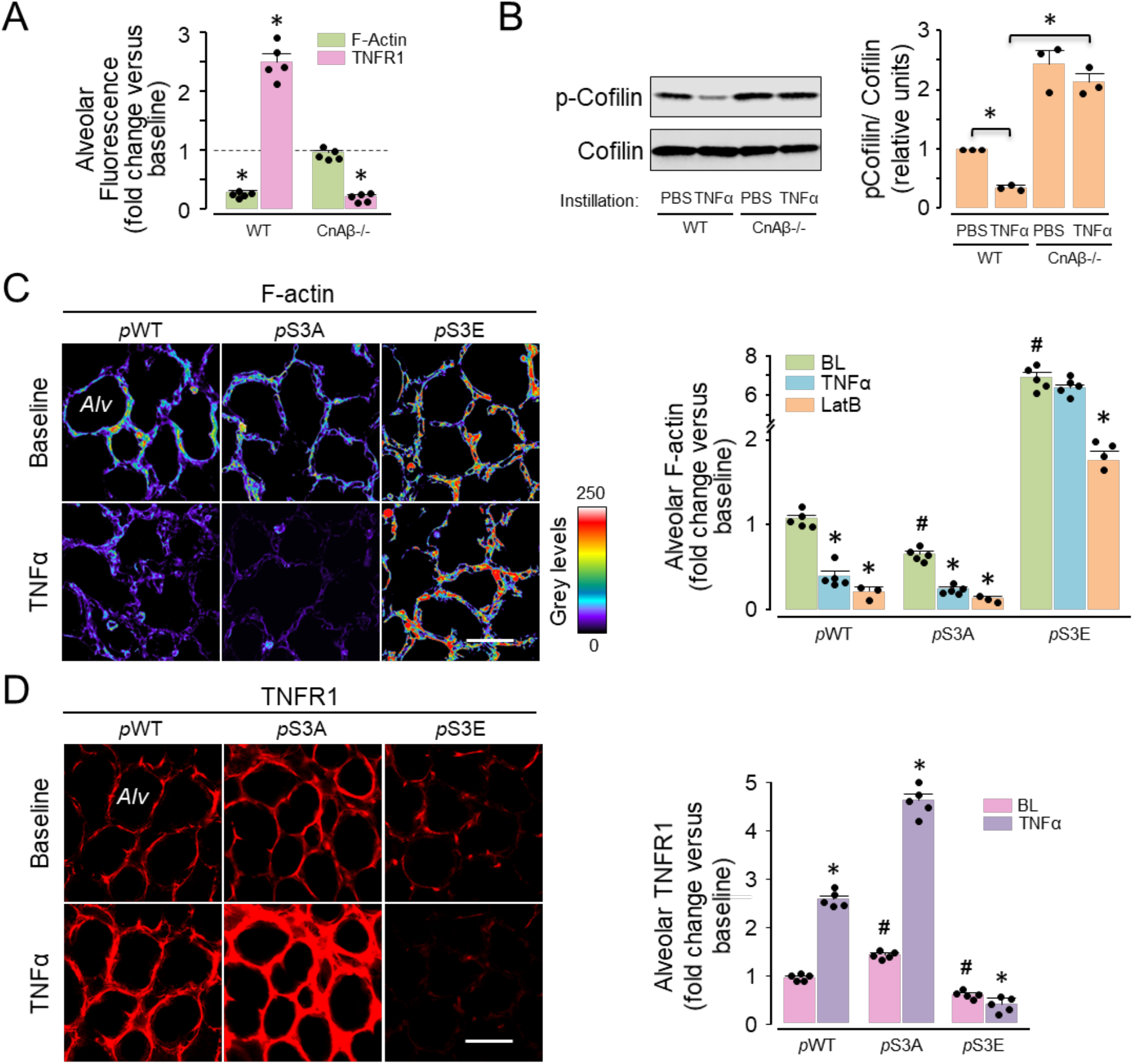
Calcineurin-dependent alveolar TNFR1 expression. (**A**) Alveoli were optically imaged to quantify fluorescence of F-actin (Lifeact) and TNFR1 (immunofluorescence) in wild type (*WT*) and calcineurin-Aβ-null (*CnAβ−/−*) mice. Data were obtained as whole-image fluorescence (grey levels above background) at baseline (*dashed line*) and 30 min after alveolar microinfusion of TNFα. Microinjections of anti-TNFR1 Ab for TNFR1 detection were given prior to and 30 min after TNFα. Mean±SE, *n*=5 lungs for each group, \**p*<0.05 compared to baseline using ANOVA with Bonferroni correction. (**B**) Immunoblots and densitometry of lung lysates obtained 4 h after i.n. TNFα. Replicated 3 times, \**p*<0.05 as indicated. (**C,D**) Cofilin transfections were for wild-type plasmid (*p*WT), and for constitutively active (*p*S3A), or inactive (*p*S3E) cofilin mutants. *LatB,* latrunculin B. Images in **C** show alveolar F-actin in terms of rhodamine-phalloidin fluorescence at baseline (*upper panel)* and 30 minutes after alveolar injection of TNFα (*lower panel*). The images in **D** were obtained at baseline after the first microinjection of anti-TNFR1 Ab (*upper panel)*. A second Ab microinjection for TNFR1 detection was given 30 min after TNFα microinjection (*lower panel)*). The bars in **C** and **D** are quantifications of whole-image fluorescence (mean±SE). *Alv,* alveolus; scale bars, 50 μm. Each dot shows data for a single lung. *n=*5 for all groups, except LatB (*n*=3 for *p*WT and *p*S3A, *n*=4 for *p*S3E). *p*<0.05 vs baseline for same group (*) using 2-tailed *t* test, or baseline for the *p*WT group (^**#**^) using ANOVA with Bonferroni correction.

To further explore the role of cofilin, we transfected the alveolar epithelium with plasmids to express wild type cofilin (*p*WT), or cofilin mutants that cannot be phosphorylated (*p*S3A), or are constitutively phosphorylated (*p*S3E). Hence these mutants are respectively, constitutively active, or inactive (Supplemental Figure 3) (25–27). As compared to *p*WT-expressing epithelium, baseline F-actin was lower in *p*S3A-, but higher in *p*S3E-expressing epithelium (Figure 2C). Alveolar TNFα microinjection decreased F-actin in *p*S3A-, but not in *p*S3E-expressing epithelium (Figure 2C). This lack of effect was not due to detection failure, since the actin depolymerizing agent, latrunculin B decreased F-actin in *p*S3E-expressing epithelium (Figure 2C). The TNFα-induced TNFR1 surge was greater in *p*S3A- than WT-expressing epithelium, but blocked in *p*S3E-expressing epithelium (Figure 2D). These findings indicate that active, namely dephosphorylated cofilin was required for the TNFα-induced TNFR1 surge. Taken together, our findings indicate that a major effect of TNFα was to activate calcineurin, which then lead to cofilin dephosphorylation. As a consequence, epithelial F-actin decreased, resulting in the TNFR1 surge.

### Delivery of exogenous of Rac1 mutants modifies the actin cytoskeleton in alveolar epithelium

The GTPase, Rac1 phosphorylates p21-activated kinase, leading to LIM-kinase dependent cofilin phosphorylation, hence F-actin stabilization (28–31). To evaluate the therapeutic potential of these mechanisms, we developed non-covalent TAT-linked conjugates with His-V12Rac1 and His-N17Rac1, which are constitutively active and inactive mutants of Rac1, respectively (32). We engineered the constructs to be unstable at low pH to enable intracellular separation of TAT from the cargo protein, hence to retain the protein in the cytosol. Accordingly, when we labelled TAT and V12Rac1 with different fluorophores and injected the construct by alveolar microinjection, we could detect rapid entry of TAT-V12Rac1 in the alveolar epithelium (Figure 3A). Subsequently, TAT fluorescence progressively decreased (Figure 3A), indicating TAT exited from the epithelial cytosol, while V12Rac1 remained. We affirmed that the spatial distribution of V12Rac1 fluorescence in the epithelium matched that of the cytosolic dye, calcein red (Supplemental Figure 4A), and that the cell-impermeable fluorescence quencher, trypan blue (TB) failed to diminish V12Rac1 fluorescence (Supplemental Figure 4 A, B).

**Figure 3.**
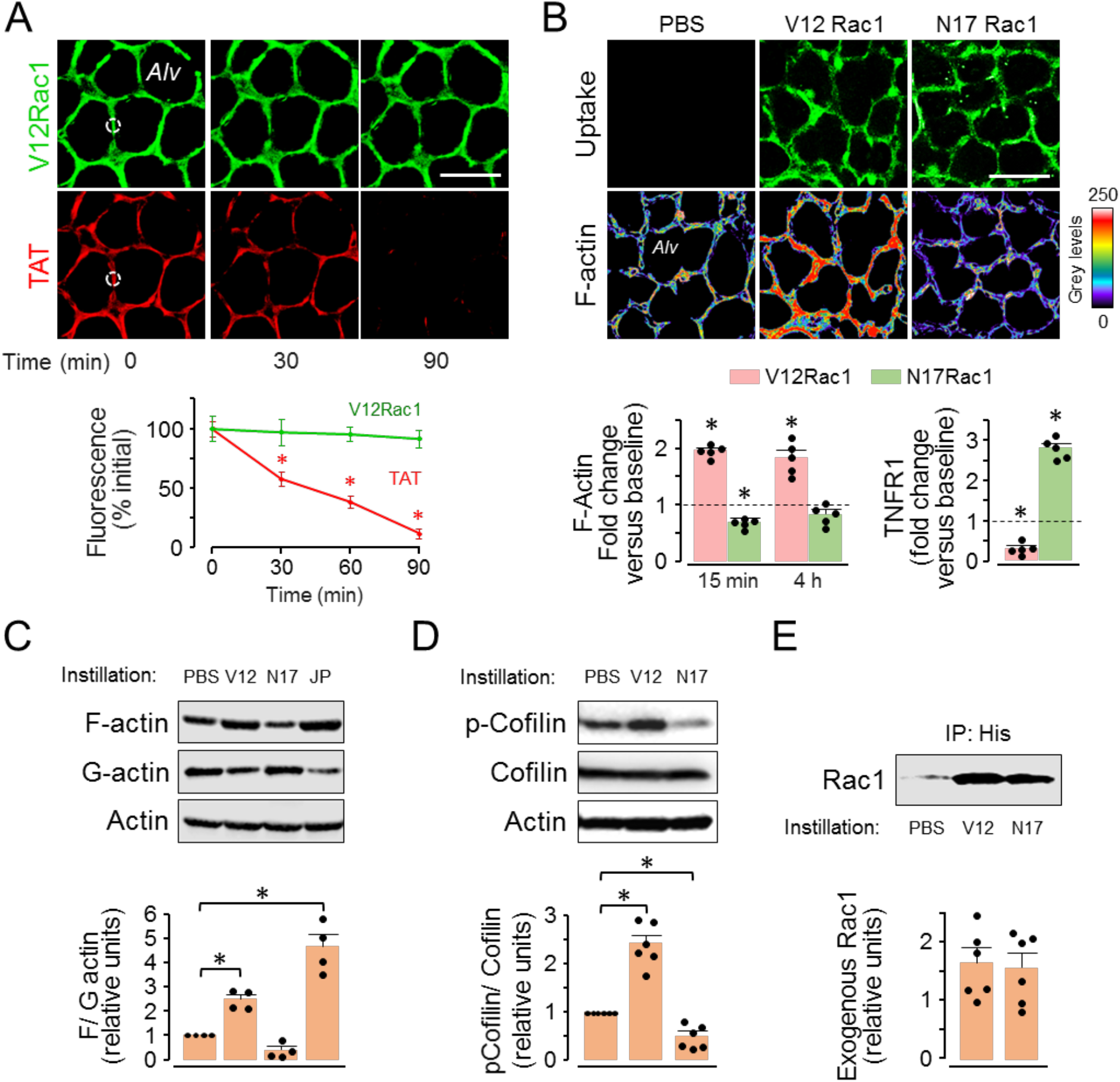
Exogenous delivery of Rac1 mutants modifies the actin cytoskeleton in alveolar epithelium. (**A**) Confocal images and the plots show changes in alveolar fluorescence of differentially tagged TAT and V12 Rac1. The tagging was done prior to conjugation. The plot shows data from ROIs (*dotted circles*) for 10 alveoli of a single lung (mean±SE). Replicated in 3 lungs. Scale bar, 50 μm. \**p*<0.05 V12Rac1 vs TAT using 2-tailed *t* test. (**B**) Images show F-actin levels 15 min after alveolar microinfusions of Rac1 mutants. Scale bars, 50 μm. Bars are whole-image quantifications of F-actin (rhodamine-phalloidin) *(left*) at indicated time points and TNFR1 expression (*right*) 30 min after TNFα microinfusion in alveoli pre-treated with TAT-Rac1 mutants. Mean±SE, *n*=5 lungs for each group. Each dot shows data for a single lung. \**p*<0.05 vs corresponding baseline (*dashed line*) using ANOVA with Bonferroni correction. (**C-E**) Immunoblots and densitometry are for lungs given indicated instillations. Tissues were harvested at 4 (**C, D**), or 24 (**E**) hours after instillations. Bands in **C** are immunoblots on lysates that were fractionated as triton-insoluble (*F-actin*), or -soluble (*G-actin*), or not fractionated (*Actin).* Bands in **E** were obtained on immunoprecipitates of His-tagged, exogenous Rac1 mutants. *PBS*, control; *V12,* TAT-V12Rac1; *N17,* TAT-N17Rac1; *JP,* jasplakinolide. All samples in blots were run simultaneously. Each blot was replicated 4 times (**C**) or 6 times (**D, E**). Each dot shows data for a single lung. \**p*<0.05 compared to PBS using ANOVA with Bonferroni correction.

To determine whether the TAT-protein constructs entered the endothelium of adjoining capillaries, we gave alveolar microinjections of TAT-V12Rac1 in which V12Rac1 was fluorophore tagged. Then, we loaded the alveolar epithelium and the endothelium with calcein red. Alveolar microinjection of the detergent, saponin eliminated epithelial, but not endothelial cytosolic fluorescence. Thus, the endothelial PM was intact and no V12Rac1 fluorescence was evident in the endothelial cytosol (Supplemental Figure 4 A, B). We conclude that the construct did not cross the alveolar barrier to enter the endothelium.

Alveolar microinjection of TAT-V12Rac1 induced rapid increase of epithelial F-actin that was sustained for at least 4 hours (Figure 3B). Although baseline F-actin was higher in AT2 than AT1 (p<0.01) (33), V12Rac1 induced a greater relative increase of F-actin in AT1 (Supplemental Figure 5). By contrast, the constitutively inactive form of Rac1, N17Rac1, which was also internalized by the epithelium, failed to increase actin (Figure 3B). The TNFα-induced TNFR1 surge was absent in TAT-V12Rac1 loaded epithelium, but present in epithelium loaded with TAT-N17Rac1 (Figure 3B). These findings indicated that epithelial loading with V12Rac1 increased F-actin, inhibiting the TNFα-induced TNFR1 surge.

To determine the extent to which these responses occurred at a whole-organ level, we carried out immunoblots on lung lysates 4 hours after i.n. instillation of the TAT-linked constructs. Our findings indicated that within 4 hours, TAT-V12Rac1, but not TAT-N17Rac1, increased F-actin, while decreasing G-actin (Figure 3C), and concomitantly increasing cofilin phosphorylation (Figure 3D). To determine longer term effects, we gave i.n. instillations of the TAT constructs. Then after 24 hours, we immunoprecipitated the Rac1 mutants from lung lysates using anti-His antibody. Both His-tagged proteins were detectable in immunoblots (Figure 3E), indicating that exogenous Rac1 mutants remained in the lung for at least 24 hours after cell internalization. Taking the imaging and the immunoblot data together, our findings indicated that epithelial loading with V12Rac1 increased F-actin, inhibiting the TNFα-induced TNFR1 surge.

### Protective effects of alveolar F-actin enhancement on ALI outcomes

To determine the role of alveolar F-actin in a mouse model of ALI, we gave i.n. LPS at lethal-dose 0 (LD0), at which there is no mouse mortality (7, 34), then determined responses after 24 hours. In optically imaged alveoli, LPS increased epithelial TNFR1 expression (Figure 4A). Concomitantly, there was alveolar inflammation, as indicated by alveolar neutrophil entry (Figure 4A). By contrast, i.n. instillation of TAT-V12Rac1 30 min prior to LPS, but not of TAT-N17Rac1, blocked the increase of TNFR1 expression, as well as the alveolar inflammation (Figure 4A). LPS induced the expected ALI responses after 24 hours, namely increased leukocytes in the bronchoalveolar lavage (BAL) (Figure 4B), increased alveolar permeability to intravascularly injected albumin (Figure 4C).

**Figure 4.**
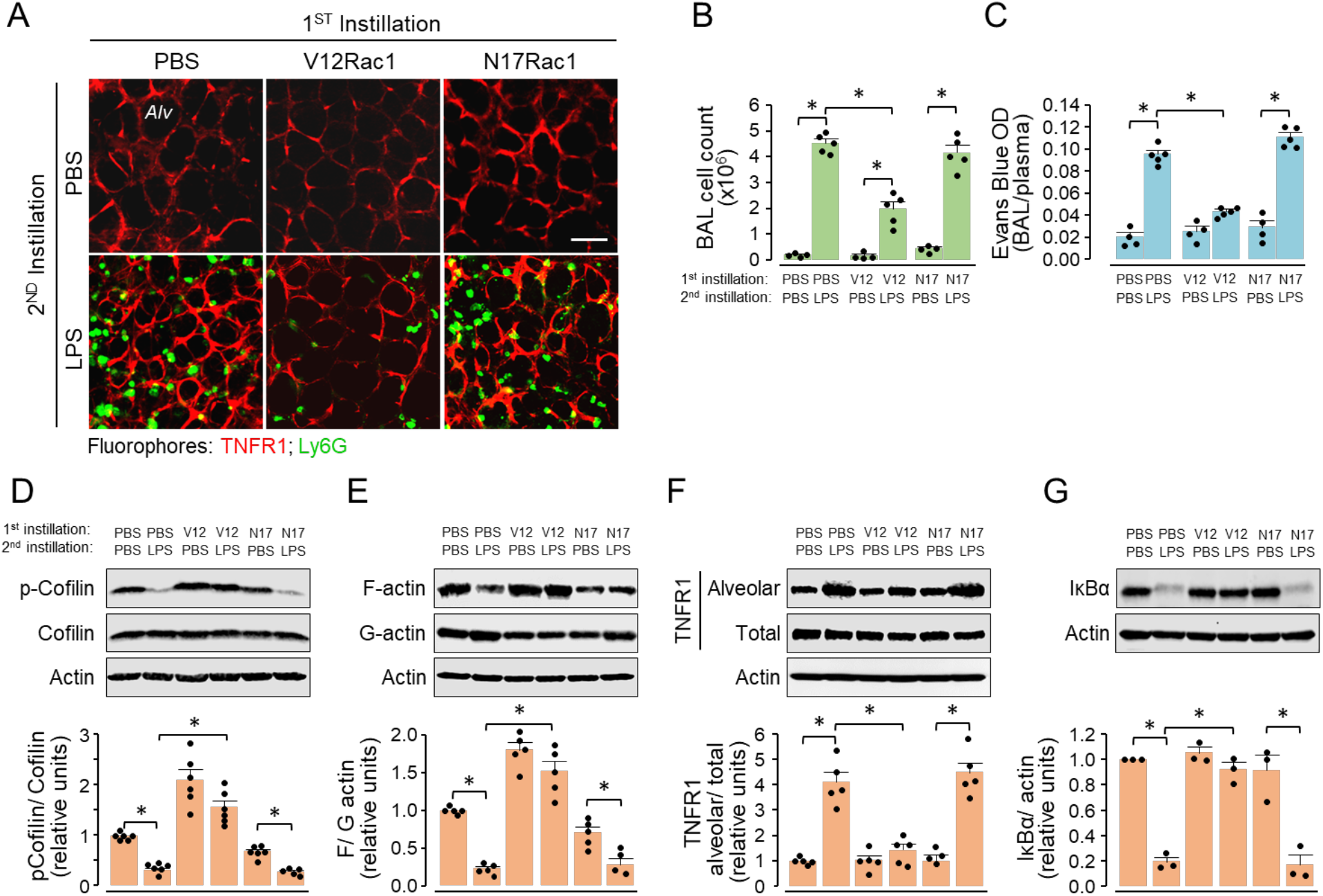
Effects of Rac1 mutants on alveolar F-actin and TNFR1 expression after LPS. Mice were given two i.n. instillations each, as indicated. Mice were separately given a 1^st^ instillation of PBS, TAT-V12Rac1, or N17-Rac1. In each mouse, a 2^nd^ instillation was given after 30 min, of PBS or LPS (sublethal dose). Data were obtained 24 h after LPS. **(A)** Confocal images show alveolar immunofluorescence of TNFR1 (red) and of neutrophils, detected by Ly6G immunofluorescence (green) after indicated treatments. Antibodies were microinjected in alveoli. Scale bar, 50 μm. *Alv*, alveolus. Replicated in 3 lungs for each group. **(B, C)** Bars show the total leukocyte count in BAL (**B**) and alveolar permeability to albumin (**C**). *OD,* optical density of Evans Blue-bound albumin. *n*=4 mice in PBS treated groups, *n*=5 mice in LPS treated groups. (**D-G**) Lung immunoblots and densitometry are for indicated proteins for total lung lysates (**D, G**), and for triton-insoluble (F-actin) and -soluble (G-actin) fractions of lung lysates (**E**). *Alveolar,* streptavidin pulldown of epithelium biotinylated *in situ*. Bars are mean±SE. Each dot shows data for a single lung. *n=*5-6 except in **G**, as indicated by the dots. \**p*<0.05 as indicated using ANOVA with Bonferroni correction.

To determine global lung responses, we carried out *in situ* biotinylation assays to determine cell-surface expression of TNFR1, as also assays of whole lung lysates for cofilin phosphorylation, F-actin and IκB. These studies indicated that LPS decreased cofilin phosphorylation (Figure 4D) as well as F-actin in 24 hours (Figure 4E) while increasing surface TNFR1 expression (Figure 4F). By contrast, pre-LPS i.n. instillation of TAT-V12Rac1, but not of TAT-N17Rac1, stabilized F-actin for 24 hours (Figure 4E) and markedly abrogated the LPS-induced enhancement of TNFR1 expression. Since NFkB activation causes alveolar inflammation, we confirmed that LPS induced IkB degradation (Figure 4G). TAT-V12Rac1, but not TAT-N17Rac1, blocked IkB degradation (Figure 4G). Since inhibition of IkB degradation inhibits NFkB activation, we interpret V12 Rac1-induced actin enhancement inhibited NFkB activation. Taking the imaging and global data together, our findings indicate that the LPS-induced proinflammatory effect of epithelial TNFR1 hyperexpression was sustained for at least 24 hours, and that these responses were inhibited by epithelial incorporation of V12Rac1.

To determine the effects of the Rac1 constructs on LPS-induced mortality, we exposed mice to a lethal LPS dose, which caused robust increases of leukocyte counts and protein concentration in the BAL (Supplemental Figure 7A), depletion of surfactant phospholipids in the BAL (Figure 5A and Supplemental Figure 7B), loss of lung compliance (Figure 5B), and ~80% mortality in 3-4 days (Figure 5C). Pretreatment with i.n.TAT-V12Rac1 30 min prior to LPS instillation protected against mortality (Supplemental Figure 6). To evaluate post-ALI therapeutic efficacy, we gave the constructs 4, or 24 hours after LPS. In the 4-hour group, V12 Rac1 mitigated LPS-induced ALI as indicated by recovery of lung compliance and BAL surfactant phospholipids (Figure 5A, B), and a 15% mortality in 3 days (Figure 5C). In the group given TAT-V12Rac1 24 hours after LPS, evaluation of ALI after a further 24 hours indicated marked reduction of lung water and BAL leukocytes (Figure 5D, E), and of mortality (Figure 5F). TAT-N17Rac1 was without effect (Figure 5F). The reduction of mortality between the 4- and 24-hour groups was not statistically significant. These findings indicate that given 30 minutes before, or 4, or 24 hours after LPS, TAT-V12Rac1 protected against ALI.

**Figure 5.**
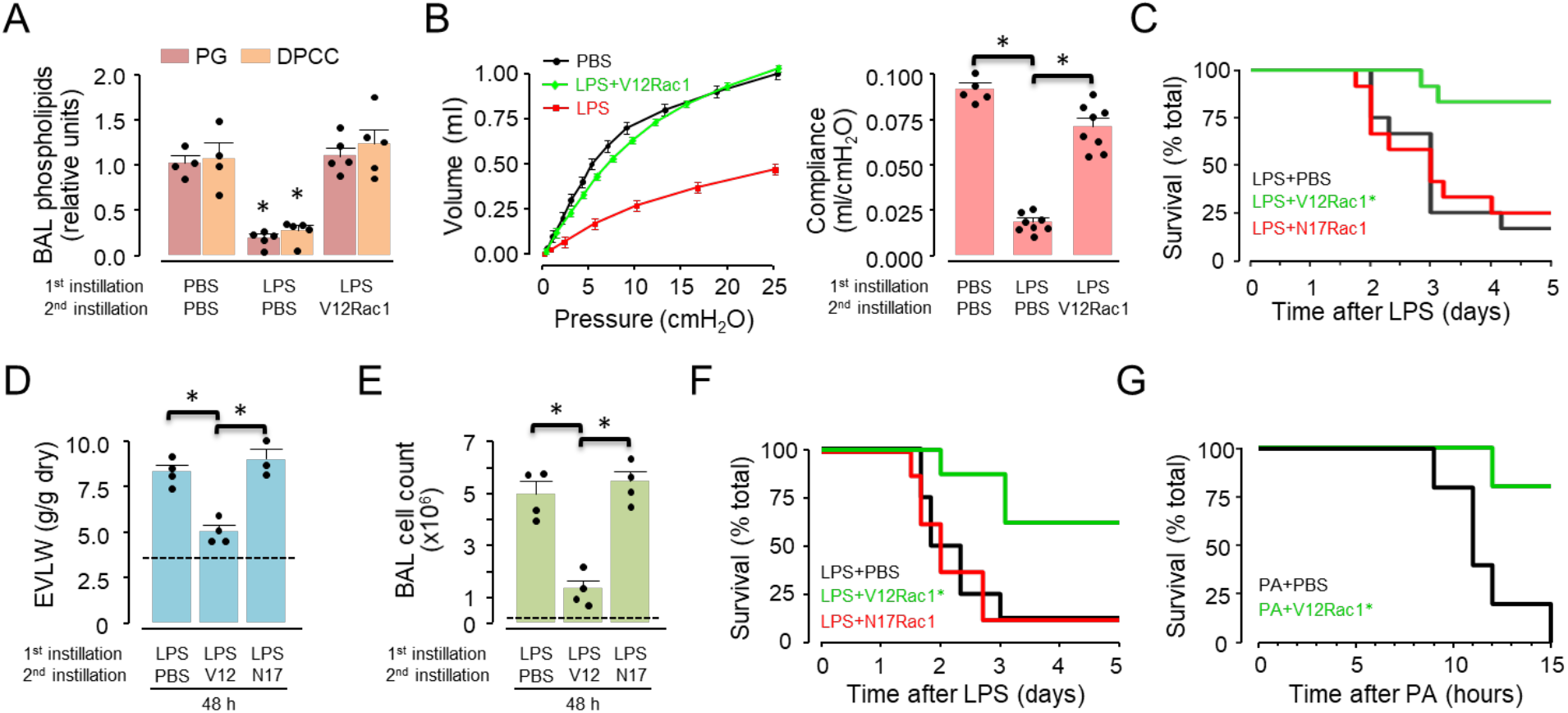
Protective effects of alveolar F-actin enhancement on ALI. (**A-F**) Mice received two i.n. instillations. The first instillation was a lethal dose of LPS. This was followed by the second instillation, which was PBS, or TAT-conjugated proteins after 4 (**A-C**), or 24 (**D-F**) hours. (**A, B**) Data were obtained 72 hours after LPS. (**A**) Bars show BAL phospholipids levels as indicated. Mean±SE, \**p*<0.05 vs. corresponding PBS group using ANOVA with Bonferroni correction. *n=*5, except PBS (*n*=4). Each dot shows data for a single lung. (**B**) Bars show lung compliance calculated from the Volume-Pressure plot. Mean±SE, \**p*<0.05 using ANOVA with Bonferroni correction. *n=*8, except PBS (*n*=5). Each dot in bar diagram shows data for a single lung. (**C**) Kaplan-Meier plots for mouse survival. For each group, *n*=12. \**p*<0.01 vs LPS+PBS using log-rank test. (**D, E**) Data were obtained 48 hours after LPS. Bars show blood-free extravascular lung water (EVLW) (**D**) and total leukocyte count in BAL (**E**). Mean±SE, \**p*<0.001 vs. as indicated using ANOVA with Bonferroni correction. *n=*4, except TAT-N17Rac1 in D (*n*=3). Each dot shows data for a single lung. Baseline values shown as dashed line (**F**) Kaplan-Meier plots for mouse survival after LPS. For each group, *n*=8. \**p*<0.01 vs LPS+PBS using log-rank test. (**G**) Kaplan-Meier plots for mouse survival after instillation of *P. aeruginosa* (PA), followed 4 hours later by TAT-Rac1 proteins. For each group, *n*= 5 \**p*<0.01 vs PA+PBS using log-rank test.

To determine whether the i.n. delivered constructs entered the capillary endothelium, we gave mice i.n. LPS at a lethal dose, or PBS, then followed 4 hours later with i.n. TAT-V12Rac1-GFP. After a further hour, we freshly isolated cells from harvested lungs, then we carried out flow cytometry analyses on the cells. These analyses indicated that V12Rac1 was taken up in the epithelium, but not the endothelium (Supplemental Figure 7C). Thus, while the imaging data indicated there was no endothelial uptake of the construct under baseline conditions (Supplemental Figure 4), the flow cytometry analyses affirm this finding and show further that endothelial uptake did not take place after LPS treatment. Taken together, our findings indicated that LPS decreased epithelial F-actin by a cofilin-mediated mechanism, increasing TNFR1 expression at the epithelial surface, augmenting lung inflammation, hence mortality. We conclude, V12Rac1 succeeded in increasing survival in LPS-induced lung injury by both prophylactic and therapeutic strategies.

LPS, an outer coat protein of Gram-negative bacteria, causes ALI by activating TLR4-induced proinflammatory signaling in pulmonary alveoli (7). However, Gram-negative bacteria such as *Pseudomonas aeruginosa* (PAO1) may induce ALI through additional mechanisms, as for example by production of exotoxin (35). To determine the efficacy of actin enhancement therapy in ALI due to live bacterial infection, we i.n. instilled PA at a dose that causes high mortality. Fifteen hours after instillation mouse mortality was 100% (Figure 5D). To determine therapeutic efficacy of i.n. TAT-V12Rac1, we administered the construct 4 hours after bacterial instillation. At this time point, lung injury was well developed as indicated by the extravascular lung water, which was two-times above baseline (Supplemental Figure 8). Despite the severe lung injury, i.n. TAT-V12Rac1 decreased mortality to ~20% at 15 h. Thus, a post-injury therapeutic strategy with i.n. TAT V12Rac1 protected against ALI caused by instillation of highly toxic bacteria.

## Discussion

We report here the first definitive evidence that TNFR1 ligation induced loss of the F-actin fence in the alveolar epithelium, causing receptor hyperexpression and alveolar injury. Rac1 delivery to the alveolar epithelium enhanced the fence, blocking the receptor hyperexpression and injury. Of translational significance, the fence enhancement protected survival in LPS- and PA-induced ALI. Together, these findings, to our knowledge, constitute the first evidence that the F-actin fence of the alveolar epithelium is a determinant of lung inflammation and injury.

We propose a sequence of events (Figure 6) in which inhaled pathogen induces macrophage-derived TNFα release. TNFR1 ligation on the alveolar epithelium increases epithelial Ca^2+^, activating the calcineurin-cofilin pathway to depolymerize F-actin. The actin fence is thus disabled, causing a surge of proinflammatory receptor expression. The F-actin depolymerizing agent, cytochalasin D also caused the receptor surge, indicating that loss of F-actin was necessary and sufficient for the effect, and that non-specific receptor-mediated mechanisms were not required. Importantly, a brief TNFα exposure induced sustained F-actin decrease in the alveolar epithelium, leading to the receptor surge, which lasted several hours, possibly corresponding to the time taken for F-actin to return to baseline levels and thereby, re-establish fence conditions. The prolonged duration of the receptor surge suggests receptor hyperexpression might be sustained during the inflammatory process if F-actin fence properties are not re-established. The extent to which this interpretation applies to other proinflammatory receptors expressed on the epithelium (2) require further investigation.

**Figure 6.**
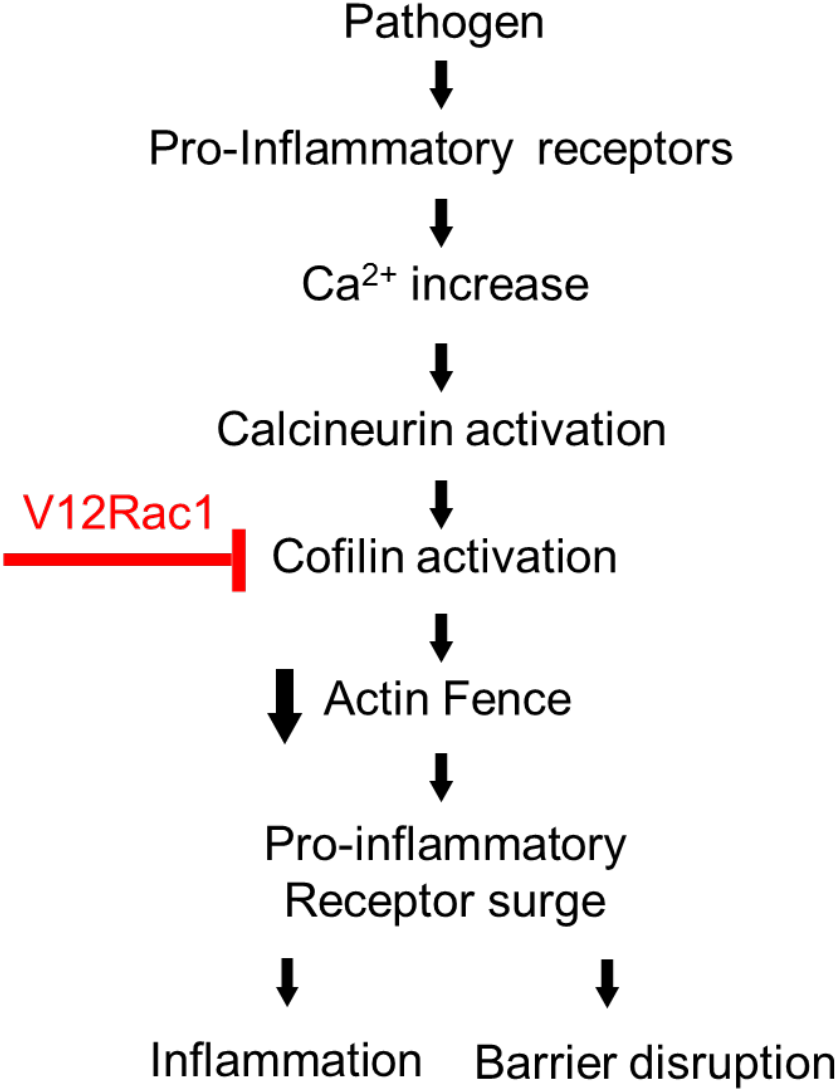
F-actin in acute lung injury. Sequence of events shows signaling mechanisms leading to F-actin-dependent proinflammatory receptor surge during Acute Lung Injury.

Actin depolymerization also destabilizes cell-junctional barriers (36), causing increased alveolar permeability to plasma proteins. Repetitions of this sequence of events may perpetuate lung injury, progressively increasing its severity and leading to the high mortality of ARDS. The fact that Rac1 delivery to the alveolar epithelium inhibited this injurious sequence indicates that actin enhancement in the epithelium was sufficient for opposing two critical players in the injury response, namely hyperexpression of pro-inflammatory receptors and weakening of the epithelial fluid barrier. Since we did not detect exogenous Rac1 in the capillary endothelium, we interpret that V12Rac1’s protective effects were predominantly due to epithelial actin enhancement.

The cell-surface expression of the receptor was AT1 restricted, with no detectable AT2 expression. TNFR1 expression on AT1 related inversely to F-actin levels, indicating that the fence was a determinant of the expression even under unstressed conditions at baseline. Cultured, AT2-like A549 cells express TNFR1 (37), although these cells do not entirely reflect properties of AT2 *in situ*. Interestingly, as we reported (33), and confirm here, F-actin is three-fold higher in AT2 than AT1, suggesting that high F-actin restricts AT2 expression of cell-surface TNFR1.

Our findings reveal novel understanding of Ca^2+^-induced F-actin regulation. An increase of the cytosolic Ca^2+^ may affect F-actin differently in different functional contexts. In the context of endothelial barrier regulation, Ca^2+^ increases lead to actin polymerization, hence stress fiber formation and barrier loss (38–40). By contrast, Ca^2+^ increases due to T cell receptor engagement at the immunological synapse induces actin depolymerization, thereby promoting the intensity and duration of T cell engagement (41). Here, the TNFα-induced Ca^2+^ transient caused actin depolymerization as evident in the rapid F-actin decrease. This response, as well as the ensuing TNFR1 surge were inhibited by blocking the Ca^2+^ transient with calcium chelator BAPTA-AM. Hence, the cytosolic Ca^2+^ is a major fence regulatory mechanism that determines TNFR1 display on the alveolar epithelium.

The Ca^2+^ effect on F-actin was mediated through the calcineurin-cofilin pathway. Several findings support this interpretation. Thus, the TNFα effects on F-actin and the surge were absent after alveolar treatment with FK-506, or in CnAβ-null mice, implicating calcineurin in the responses. We interrogated the mechanistic pathway through epithelial expressions of constitutively active, or inactive mutants of cofilin. The active mutant, which causes F-actin depolymerization, enhanced the TNFα-induced TNFR1 surge. The inactive mutant, which increased F-actin, blocked all TNFα-induced responses including the surge. These results are further evidence that loss of the actin fence was critical for epithelial hyperexpression of the proinflammatory receptor. Taken together, these findings reveal a novel sequence of fast-acting signaling events in which a receptor-mediated Ca^2+^ increase activated calcineurin, leading to cofilin-mediated loss of the F-actin fence, resulting in enhanced proinflammatory receptor display on the alveolar surface.

LPS induced the expected lung inflammation and alveolar injury, as indicated by presence of albumin leak, and significant mortality (7, 34). LPS induces secretion of multiple cytokines, including TNFα, while also increasing alveolar epithelial Ca^2+^ (7). Similar to TNFα, alveolar LPS exposure also dephosphorylated cofilin and decreased F-actin, while increasing TNFR1 expression. However importantly, all of these LPS effects were sustained for 24 hours. This finding indicates that mortality resulted from prolonged loss of the F-actin fence that caused prolonged TNFR1 hyperexpression, exacerbating alveolar inflammation. We tested this possibility by means of a CPP strategy where the plan was to increase the actin fence through intracellular delivery of V12Rac1 in order to phosphorylate, hence inactivate cofilin.

Our strategy was highly successful. Whether administered by alveolar microinjection, or by the i.n. route, the TAT-V12Rac1 construct was rapidly internalized by the alveolar epithelium. The pH-sensitive, non-covalently conjugated construct hydrolyzed intracellularly, enabling TAT diffusion out of the cell. Hence, although the protein cargo was successfully taken up by the epithelium, TAT, which contains a nuclear-localizing sequence (42) and may therefore induce unwanted transcriptional effects, was eliminated. We consider this TAT elimination from the alveolar epithelium a strength of our CPP strategy, since such an elimination would abrogate the likelihood of transcription-induced long-term toxicity due to TAT. Although reports indicate that extracellular TAT is rapidly removed from the lung (43, 44), further studies are required to understand mechanisms underlying the removal process.

The instilled TAT-V12Rac1 construct was not detectable in the capillary endothelium before or after induction of alveolar inflammation. Thus, once dissociated from TAT, non-conjugated V12Rac1 protein remained confined to the epithelial cytosol and did not enter endothelial cells. We affirmed this result by assaying the membrane permeabilizing effects of saponin on cell fluorescence. Given by alveolar injection, saponin removed epithelial, but not endothelial fluorescence (sFig. 4), indicating that the construct was localized only to the alveolar compartment. This conclusion is further supported by our flow cytometry results showing that even after LPS treatment, a condition that increased alveolar permeability, there was no endothelial uptake of V12Rac1 (SFig. 8C). Therefore, TAT-V12Rac1 did not modify endothelial F-actin. We consider it unlikely that it directly affected endothelial barrier determinants, such as focal adhesions and junctional proteins (45–47).

Following delivery of the CPP, we could detect V12Rac1 in the epithelium for at least 24 hours, during which epithelial F-actin also remained elevated and the TNFR1 hyperexpression was abrogated. These findings show for the first time, that constitutively active Rac1 increases F-actin *in vivo*, and that the induced F-actin remains stable for sufficient durations to be therapeutically effective. Importantly, our findings indicate that pre-treatment with i.n. TAT-V12Rac1 markedly reduced mortality due to LPS. We point out however, that while our studies support a protective effect of the actin fence in the context of TNFR1 expression, the extent to which actin fence strengthening modifies the effects of other ALI-relevant receptors, such as the proinflammatory interleukin1β receptor, or the anti-inflammatory TNFR2 (48) requires further study. Nevertheless, our findings indicate that pre-treatment strategy with V12Rac1 might be a feasible prophylactic option against ALI.

To determine the therapeutic efficacy of TAT-V12Rac1 as a curative agent, we instilled the construct under conditions of established pathology, namely 4 or 24 hours after instilling a lethal LPS dose that caused major alveolar hyperpermeability and inflammation. These post-LPS V12Rac1instillations were also protective against mortality. An important conclusion is that fence enhancement in the alveolar epithelium mitigates ALI well into the progressive phase of the disease. However, since the mortality protection was less for the delayed intervention group, fence enhancement therapy might be more effective in early than late stages of ALI.

We also considered that infection by live bacteria may induce injury mechanisms that are more extensive than those due to LPS alone. Accordingly, we determined the effects of the construct in the presence of lung infection by a highly lethal dose of PA, an organism that is a major cause of ALI and ARDS (49). In wild-type mice, this dose of PA induced catastrophic mortality in 15 hours. However, when we gave TAT-V12Rac1 after inducing infection, the mortality was markedly abrogated. These findings add to the translational significance of our study, which reveals the therapeutic strength of V12Rac1 in the setting of ongoing lung inflammation.

Antibody inhibition of TNFR1 has been advanced as a possible therapeutic strategy for ALI (50, 51). However, antibody therapy was protective for ALI not associated with mortality (50, 51). Hence, these reported findings are not directly comparable to the present mortality-causing inflammatory responses, which could be encountered in clinical ARDS. In severe inflammation, cell-directed therapy as we propose, may be more effective.

Since Rac1 plays a role in multiple cellular processes (52–54), we considered whether its sustained epithelial presence affected alveolar function. We approached this issue by quantifying lung surfactant secretion and lung compliance, metrices that report adequacy of overall alveolar homeostasis. Maintenance of surfactant secretion reflects adequacy of several epithelial regulatory parameters, such as cytosolic Ca^2+^ regulation (55). The fact that V12Rac1 protected these critical functional responses despite ALI, indicates that V12Rac1 treatment not only did not interfere with major aspects of epithelial function, but that the treatment reinstated alveolar homeostasis in ALI.

Although our study indicates that there were no overall negative consequences attributable to V12Rac1, to support clinical application further data are required to clarify issues including the dynamics of elimination of the internalized V12Rac1, as well as protective efficacy in large animal models of ALI. Since pharmacologic therapy for ALI remains elusive, and since the therapeutic efficacy of cellular delivery of protein constructs has not been previously evaluated, we conclude that our novel V12Rac1 construct may have potential as therapeutic strategy for ALI, as well as for severe inflammatory diseases in other organs.

## Materials and Methods

### Reagents

The following reagents were purchased: fluorophores calcein AM (5–10 μM), calcein red-orange AM (5 μM), lysotracker red (LTR; 100nM) and Fluo-4 (10 μM) (Life Technologies), and Alexa-488-phalloidin and Rhodamine-phalloidin (Invitrogen). Human recombinant TNFα (10-100 ng/ml, BD Biosciences), BAPTA-AM (100 μM, Life Technologies). FK506 (100μM), LPS (1-10 mg/kg), cytocholasin D (100nM), jasplakinolide (100 nM), latrunculin B (50 nM), trypan blue (0.01%), DTT (1 mM), Turk’s solution and trypan blue were purchased (Sigma-Aldrich). Protein A/G-agarose beads (Santa Cruz). Saponin (Calbiochem). EZ-Link *N*-hydroxysuccinimide-SS-biotin (#21331) and streptavidin-Sepharose beads (#20357) (Thermo Fisher Scientific).

### Antibodies

mAb MCA2350 against the TNFR1 extracellular epitope was purchased (AbD Serotec). Antibodies were fluorescently labeled with Alexa Fluor 633 using standard protocols. Antibody against GFP (sc-9996) and IκBα antibody (sc-371-G) were from Santa Cruz Biotechnology. For immunoblotting experiments, we purchased Abs against p-Cofilin (# 3313) and Cofilin (# 5175) from Cell Signaling. Antibody aganist TNFR1 (sc-8436), His-probe (sc-8036) and Rac1 (sc-6084) were from Santa Cruz Biotechnology. Anti-actin Ab (# 2066) was purchased from Sigma. Fluorescence-tagged Abs against mouse CD45 (# 103126) were from BioLegend and anti-mouse Ly6G (#11-9668), CD31 (#12-0311) and T1α (#25-5381) were from eBiosciences.

### Animals

The Institutional Animal Care and Use Committee of Columbia University Medical Center approved all animal procedures. Pathogen-free, adult male wild type Swiss Webster mice (6–10 weeks old) were purchased from Taconic Biosciences. Calcineurin Aβ-null mice were provided by Dr. Jeffery D. Molkentin (Divisions of Molecular Cardiovascular Biology Children’s Hospital Medical Center, Cincinnati, OH, USA). Age, gender and strain-matched WT mice were purchased from Jackson Laboratory. For mouse anesthesia for i.n. instillations and surgical procedures, we gave inhaled isoflurane (3%) and i.p. injections of ketamine and xylazine (100 mg/kg and 5 mg/kg respectively).

### Isolated, blood-perfused lungs

Lungs were prepared according to our reported method (7, 34, 56). Briefly, we excised intact lungs from anesthetized mice. Lungs were inflated through a tracheal cannula with a gas mixture (30% O_2_, 6% CO_2_, balance N_2_) and continuously pump-perfused with autologous blood (final hematocrit, 10%) through cannulas in the pulmonary artery and left atrium at constant flow rate of 0.4–0.5 ml/min at 37°C. Blood was diluted in HEPES buffer (150 mM Na^+^, 5 mM K^+^, 1.0 mM Ca^2+^, 1 mM Mg^2+^, and 20 mM HEPES at pH 7.4) supplemented with 4% dextran (70 kDa; TCI America) and 1% FBS (HyClone, Thermo Fisher Scientific) at osmolarity 295 mosM (Fiske Micro-Osmometer, Fiske Associates). Airway, pulmonary artery and left atrial pressures were held constant at 5, 10 and 3 cm H_2_O, respectively during microscopy.

### Alveolar microinjection and *in situ* immunofluorescence

We used isolated blood-perfused mouse lungs for imaging experiments. To load the alveolar epithelium with fluorophores, reagents, solutions, and antibodies, we micropunctured single alveoli with hand-beveled glass micropipettes (opening diameter 5-8 μm) under bright-field microscopy and microinstilled alveoli with solutions. Instilled solutions spread from the micropunctured alveolus to at least 15 neighboring alveoli within lung acini (57, 58). To carry out live immunofluorescence, alveolar antibody infusions were followed by washout with PBS. To assess fluorophore internalization, microinfusion of fluorophore-conjugated agents was followed by alveolar infusion of the membrane-impermeable fluorescence quencher trypan blue (0.01% solution, 10 minutes) (9).

To carry out *in situ* rhodamine-phalloidin and anti-GFP immunofluorescence alveoli were fixed by continuous microinfusion of 4% paraformaldehyde (15 minutes) and permeabilized by infusion of 0.1% triton X-100 (5 minutes). Then, alveoli were microinfused with fluorescence (Alexa Fluor 488 or rhodamine)-conjugated phalloidin or antibody for 10 minutes. For antibody staining we blocked the tissue for 20 min with 1% fetal bovine serum (FBS) solution, then microinfused primary antibody (30 min) and secondary antibody (30 min) in 1 % FBS containing Hepes buffer. Unbound agents were removed by 10 min washout with Hepes buffer containing 1% FBS and 0.01% Tween 20.

### Live alveolar imaging

We imaged alveoli of isolated, blood-perfused lungs, held at constant inflation, by laser scanning confocal microscopy (LSM 510; Carl Zeiss Microscopy, Heidelberg, Germany) using a ×40 water immersion objective (numerical aperture 0.80, Achroplan; Carl Zeiss Microscopy) per our reported protocol (7, 34, 56). During concurrent application of multiple fluorophores, we confirmed absence of bleed-through between different fluorescence emission channels. As indicated in the Figure Legends, fluorescence was quantified across a grey level range of 10-255 for specified regions of interest (ROIs), or for the entire image (MetaMorph 7.8, Universal Imaging). To eliminate background fluorescence, the fluorescence gain was set to give images with zero grey levels in the absence of added fluorophore. Laser power, gain and optical thickness were held constant across experiments for each fluorophore.

In case of an excessive fluorescence response, we reduced gain to prevent fluorescence saturation, then corrected grey levels against the linear relationship between grey levels and gain that we determined for each fluorophore.

We imaged TNFR1 expression as immunofluorescence of an ectodomain-specific fluorescent Ab (monoclonal antibody MCA2350; AbD Serotec, Raleigh, NC, USA) (9), which we gave by alveolar micropuncture. To image F-actin expression, in live tissue we detected fluorescence of a transfected F-actin probe (Lifeact-RFP; gift of Dr. Roland Wedlich-Soldner, Max Planck Institute, Martinsried, Germany) (33, 59), or in fixed and permeabilized tissue by rhodamine-phalloidin fluorescence. For Ca^2+^ imaging, we loaded the alveolar epithelium with the Ca^2+^ probe, fluo-4 (Invitrogen). Ca^2+^ and Lifeact images were acquired at 1 image/10 seconds.

### *In vivo* transfection

Lifeact-RFP construct was a gift of Dr. Roland Wedlich-Soldner (Max Planck Institute, Martinsried, Germany). Plasmids for wild-type cofilin (*p*WT), an activated mutant (*p*S3A), and an inactive mutant (*p*S3E) were generously provided by Dr. Stuart S. Martin (Department of Physiology, University of Maryland School of Medicine, Baltimore, MD, USA) (26). Using our established methods(7, 34, 56), we complexed plasmid DNA (75μg) with freshly extruded unilamellar liposomes (20 μg/μl; 100 nm pore size; DOTAP, Avanti Lipids) in sterile Opti-MEM (Invitrogen). We administered plasmid DNA-liposomes mix by intranasal instillation in anesthetized mice. Imaging experiments were carried out 48 hours after transfection.

### TAT-Rac1 conjugation

We cloned cDNA of V12 Rac1 and N17Rac1 (gift of Dr. P. Jurdic (Laboratoire de Biologie Moléculaire et Cellulaire, Ecole Normale Supérieure de Lyon, Lyon, France)) into a histidine (His) transfer vector. Then to make recombinant virus, we co-transfected Sf9 cells with the cloned cDNA and viral DNA (BaculoGold). We infected Sf9 cells with the recombinant virus to express and purify the His-tagged proteins using a nickel-nitrilotriacetic acid (Ni-NTA, Qiagen) column. For non-covalent, pH-dependent conjugation of TAT peptide to the Rac1 proteins, we synthesized nitrilotriacetic acid (NTA) to the N-terminus of TAT (amino acids 48-60, CHI Scientific, Maynard, Mass.). Then, we combined equimolar amounts of NTA-TAT (500 μM in 20 μl sterile, calcium-, magnesium-free PBS) and copper sulfate (500 μM in 20 μl deionized water) for 30 minutes at room temperature, to obtain copper-containing NTA-TAT. Purified His-Rac1 (V12 Rac1 or N17 Rac1) was dissolved in dipositive cation free PBS (20 μl) and then added to the NTA-TAT-Cu mixture. The reaction mixture was left at room temperature for 90 min. In this final mixture, component volumes were adjusted to establish concentrations of NTA-TAT at 50 μM and of His-Rac1 at 500 μg/kg.

### ALI

Lung injury was induced by i.n. instillation of LPS in sterile phosphate-buffered saline (PBS), or of live *Pseudomonas aeruginosa (strain* O1). LPS concentration was 1 mg/kg (non-lethal dose) for ALI experiments and 10 mg/kg (lethal dose) for the survival experiments (7, 33). PAO1 was given at a concentration of 2.5 ×10^7^ colony-forming units [cfu] in 50 μl of PBS. TAT-Rac1V12 or TAT-Rac1 N17 (0.5 mg/kg in 40 μl PBS) were given by i.n. instillation 30 minutes before or 4 hours after the induction of injury. For control, we instilled an equal volume of sterile PBS. Mice were anesthetized during the instillations.

### Bacterial preparation

*Pseudomonas aeruginosa strain PAO1* were grown on Luria–Bertani (LB; MP Biomedicals) agar at 37 °C. We inoculated LB broth with a single colony, then grew the bacteria overnight in a shaking incubator at 37°C and 250 rpm (Innova42, New Brunswick Scientific). We added 100 μl of the overnight broth culture to 10 ml fresh LB broth and propagated the bacteria in a shaking incubator to optical density (OD) of 0.5 at 600 nm (SPECTRAmax Plus, Molecular Devices). For infection we centrifuged and re-suspended 1 ml of culture (OD_600nm_ = 0.5) in 1ml of PBS, then administered 50 μl of bacterial suspension to deliver 2.5×10^7^ cfu per mouse.

### Analysis of Bronchoalveolar (BAL) fluid

The lungs of anesthetized mice were lavage 5 times with ice-cold PBS without calcium and magnesium (1 ml each time) through a tracheal cannula. Collected BAL fluids were centrifuged for 10 minutes at 350*g* and 4°C. For cell counts, pellets were re-suspended in 1 ml of PBS and stained with Turk’s solution. Cell counts were determined using a hemocytometer (Hausser Scientific). The supernatant was analyzed for protein concentration. For phospholipid analyses, supernatants were centrifuged (27,000*g*, 60 min, 4°C) to obtain a pellet containing the “large aggregate” (LA) fraction of alveolar surfactant (60–62). The LA fraction was resuspended in sterile PBS and its phospholipid composition was analyzed by thin-layer chromatography (TLC) using standard procedures following lipid extraction from the LA fraction (60, 61). Briefly, 2:1 (v/v) chloroform:methanol mixture was added to resuspended LA, then the lipid rich chloroform phase was recovered after vortexing and centrifugation. The chloroform was evaporated to dryness under N_2_. The lipid extract was reconstituted in 50 μl acetonitrile and applied to a TLC plate. Lipids were separated employing a mixture of 65:25:4 (v/v/v) chlorophorm:methanol:water as a mobile phase followed by lipid visualization in an iodine chamber.

### Evaluation of Acute Lung Injury

We followed reported protocols for quantifications of alveolar permeability (34) and extravascular lung water (EVLW) (56, 63). Briefly, alveolar permeability was quantified as the BAL-plasma ratio of Evans Blue-albumin (EB-albumin) concentrations, 4 hours after i.v. EB-albumin. EVLW was the wet-dry ratio of the lung homogenate, corrected for blood water content. To quantify lung compliance (CL), mouse lungs were inflated to 25 cm H_2_O airway pressure (P). Then, decreases in P were recorded for step decreases in lung volume (V). CL was calculated as the slope of the linear regression of V against P for P=0-10 cmH_2_O. Data for each lung are means of two replicated V-P plots.

### Survival assessment

Animals were randomly divided for the experiments groups. Anesthetized mice were treated with intranasal instillation of TAT-Rac1 conjugates (0.5 mg/kg) 30 minutes before or 4 hours after intranasal instillation of LPS at a lethal dose (10 mg/kg). The mice were scored at frequent intervals following LPS administration, in agreement with the approved Animal Care Protocol. At each assessment, the animals were scores by a blinded investigator applying scoring system which includes measuring of body weight, evaluation of respiration, activity and grooming.

### Lung cell isolation and flow cytometry

To isolate lung cells, we buffer perfused lungs through vascular cannulas to clear blood, and then minced and passed the tissue through 40 μm cell strainers (BD Biosciences) to obtained single cell suspension. For flow cytometry, we surface stained the cells by incubating the suspension with fluorophore-conjugated Abs (15 min, 4 °C). Alveolar epithelial cells were identified as CD45^-^CD31^-^ T1α ^+^ and endothelial cells were identified as CD45^-^CD31^+^T1α^-^ We analyzed cells by flow cytometry (LSR II, BD Biosciences) and using standard software (FlowJO, Tree Star Inc.).

### Immunoprecipitation and Western blot analysis

Lungs were cleared from BAL leukocytes and blood by repeated lavage via tracheal cannula and vascular perfusion with 10 ml of ice cold PBS respectively. Freshly dissected lungs were homogenized (Tissue Tearor; Biospec Products) in Pierce RIPA buffer (#89901, Thermo Fisher Scientific) containing Halt protease and phosphatase inhibitor cocktail (#78442, Thermo Scientific). The lysates were cleared by centrifugation for 20 min at 16,000 *g*. Protein concentrations were determined using a commercial kit (Pierce BCA Protein Assay Kit, Thermo Fisher Scientific), BSA standards (2 mg/ml in 0.9% saline; Thermo Fisher Scientific) and a plate reader (SPECTRAmax Plus, Molecular Devices). Samples containing equal amounts (75-100 μg) of proteins were resuspended in Laemmli sample buffer, boiled for 5 min, separated by SDS-PAGE and electrotransferred onto nitrocellulose membrane (# 1620115, Bio-Rad) overnight at 4°C.

Exogenous Rac1 mutant proteins were immunoprecipitated from equal amounts of whole lung lysates (1mg) with anti-His antibody (1 μg) overnight at 4°C and protein A/G PLUS-agarose beads (20 μl) were added to the samples and incubated for additional 4 hours at 4°C. The beads were washed five times with lysis buffer, resuspended in 2× Laemmli sample buffer, and analyzed by SDS-PAGE and Western blotting. We used primary and secondary antibodies following the manufacturer’s instructions. We imaged the blots using a Kodak molecular imaging station (IS4000MM) and Odyssey Fc Imaging System (LI-COR Biosciences). Quantification of protein levels was performed by densitometry scanning with ImageJ (NIH). Values were normalized to the PBS lane from the same experimental set.

### Biotinylation of alveolar surface protein expression

The lungs of anesthetized mice were lavaged 3 times with ice-cold PBS to remove leucocytes from the airspace. Alveolar surface proteins were labeled for 20 min using 1 mg/ml EZ-Link *N*-hydroxysuccinimide-SS-biotin instilled through a tracheal cannula, followed by repeated lavage with PBS containing 50 mM glycine to quench unbound biotin. Lung tissue were lysed in modified RIPA buffer (50 mM Tris-HCl, pH 8, 150 mM NaCl, 1% NP-40, 1% sodium deoxycholate, and protease inhibitors). Biotinylated proteins were pulled down with streptavidin beads overnight at 4°C on a rocking platform from 500μg lungs lysates and analyzed by sodium dodecyl sulfate-polyacrylamide gel electrophoresis (SDS-PAGE) and Western blotting.

### Lung fractioning for G-actin and F-actin

A two-step solubilization was carried out by our reported method (36). Briefly, lung tissue cleared from blood and BAL leukocytes were homogenized and solubilized in 400 μl of mild lysis buffer (50 mM NaCl, 10 mM Pipes, pH 6.8, 3 mM MgCl2, 0.05% Triton X-100, 300 mM sucrose) for 20 min at 4 °C on a rocking platform. The whole lung lysates were clarified by centrifugation at 10,000 *g* for 10 min. The supernatants, Triton-X 100 soluble fraction, were collected and used to determine G-actin. The resulting pellet was re-suspended in 200 μl of stronger solubilizing buffer (15 mM Tris, pH 7.5, 5 mM EDTA, 2.5 mM EGTA, 1% SDS) and incubated at 95° C for 10 min to dissolve the pellet and then volume was adjusted to 400 μl with RIPA buffer. The resulting lysates were clarified by centrifugation at 16,000 *g* for 10 min. Collected supernatant, Triton-X 100 insoluble fraction were used to determine F-actin. Extracted proteins (15μg) were subjected to SDS-PAGE and Western blotting analysis.

### Data analysis

Group numbers were designed to enable detection of statistically significant differences with a power of 85%. For imaging experiments carried out with a paired protocol, baseline and test conditions were obtained in the same alveoli and at least 5 determinations were obtained for each lung. These determinations were averaged to obtain a mean for each condition in each lung. There was no statistically significant within-lung variability of effect size. The means for each lung were pooled for the group to obtain mean± SEM, where *n* represents the number of lungs. The per-lung means are shown in the bar diagrams. Group means were compared by ANOVA with Bonferroni correction. Data for the same alveolar segment were compared by the paired t-test. Survival rates were analyzed by the Kaplan-Meyer log rank test. We accepted significance at p<0.05.

## Author contributions

G.A.G. deigned, carried out and analyzed all experiments; S.R.D. and G.J. carried out the recombinant protein purification and conjugation; M.N.I. contributed to the imaging experiments and survival studies; K.W. contributed to calcium imaging experiments; I.O.S. contributed to the TLC experiments; L.L. contributed to the EVLW experiments; S.B. contributed to the plan; J.B. designed the overall project and wrote the final manuscript. All authors contributed to the writing.

## Acknowledgments

M. Wei assisted with Rac1 protein expression.

---

**Supplemental Figure 1.**
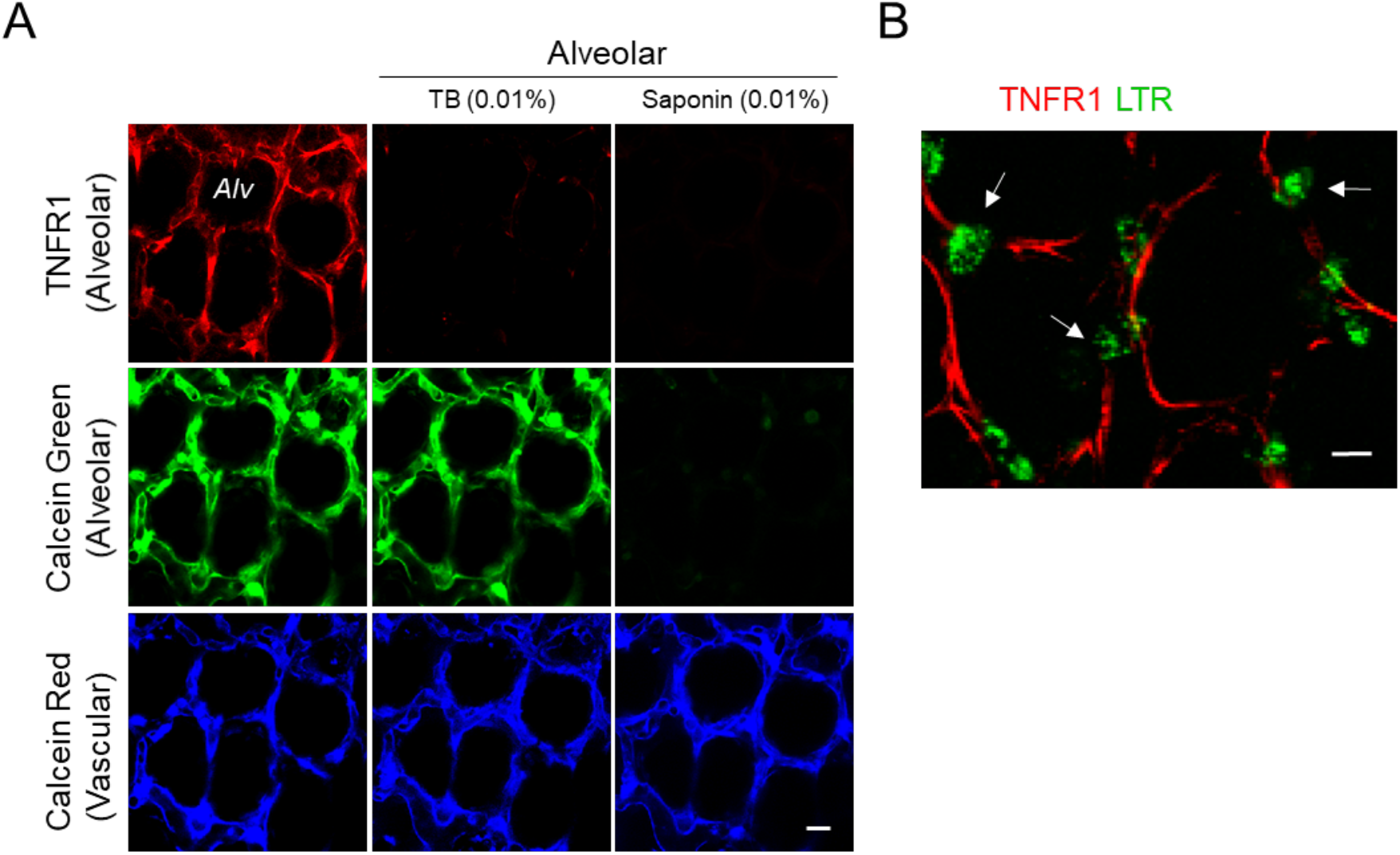
Alveolar expression of TNFR1. **(A)** Confocal images show live alveolar epithelial (top and middle row) and subjacent vascular endothelial fluorescence (bottom row) at baseline (first column), and following trypan blue (second column) and saponin (third column) treatments. In the top and middle panels, alveolar microinjections of anti-TNFR1 mAb and calcein green label the same alveoli. The bottom panel shows microvascular endothelium loaded with vascular infusion of calcein red (blue pseudocolor). Alveoli were microinfused with the membrane-impermeable fluorescence quencher trypan blue (TB) and the membrane permeabilizing agent, saponin as indicated. Note, that TB abolished TNFR1 fluorescence, indicating that the immunofluorescence was on the epithelial surface, and saponin caused fluorescence loss of epithelial, but not of endothelial cytosolic fluorescence, indicating that alveolar injections targeted only the alveolar epithelium. (**B**) Live image of alveoli shows fluorescence of TNFR1 (red). AT2 cells (*arrows*) are identified by Lysotracker fluorescence (green). Note, AT2 cells do not express TNFR1 fluorescence. Scale bar, 10 μm. Replicated in 5 lungs.

**Supplemental Figure 2.**
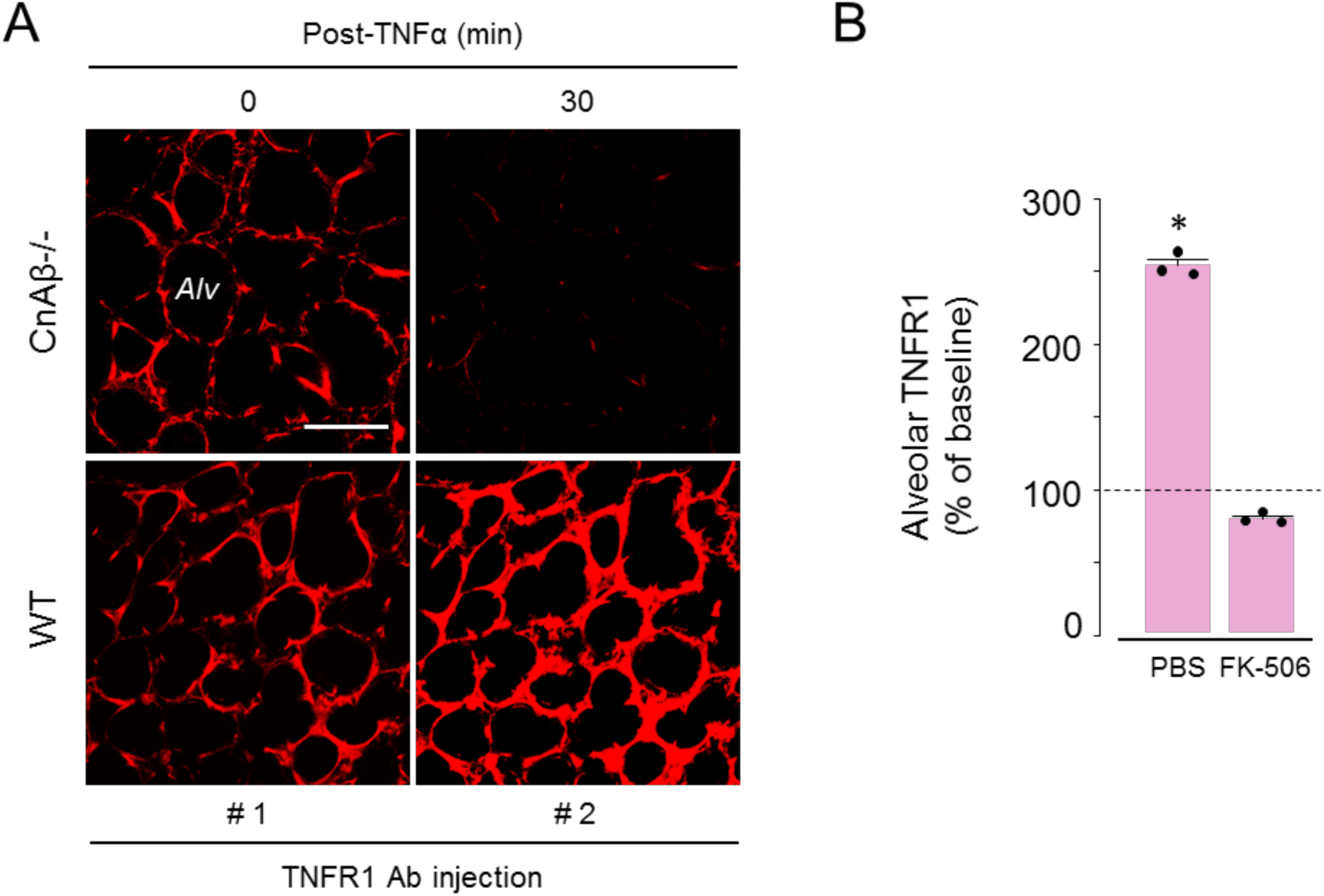
Calcineurin determines alveolar TNFR1 expression. (**A**) Confocal images show alveolar TNFR1 expression before and after alveolar TNFα microinjection in wild type (*WT*) and calcineurin-Aβ null (*CnAβ−/−*) mice. Note, 30-min images show recovery of TNFR1 expression in *WT* but not in *CnAβ* −/−. *Alv*, alveolus. Scale bars, 50 μm. Replicated in 3 lungs. (**B**) Data were obtained as whole-image fluorescence at baseline (*dashed line*) and 30 min after alveolar microinfusion of TNFα in lungs pre-treated with *FK-506*. Microinjections of anti-TNFR1 Ab for TNFR1 detection were given prior to and 30 min after TNFα. Mean±SE, *n*=3 lungs for each group, \**p*<0.05 vs baseline using 2-tailed *t* test. Each dot shows data for a single lung.

**Supplemental Figure 3.**
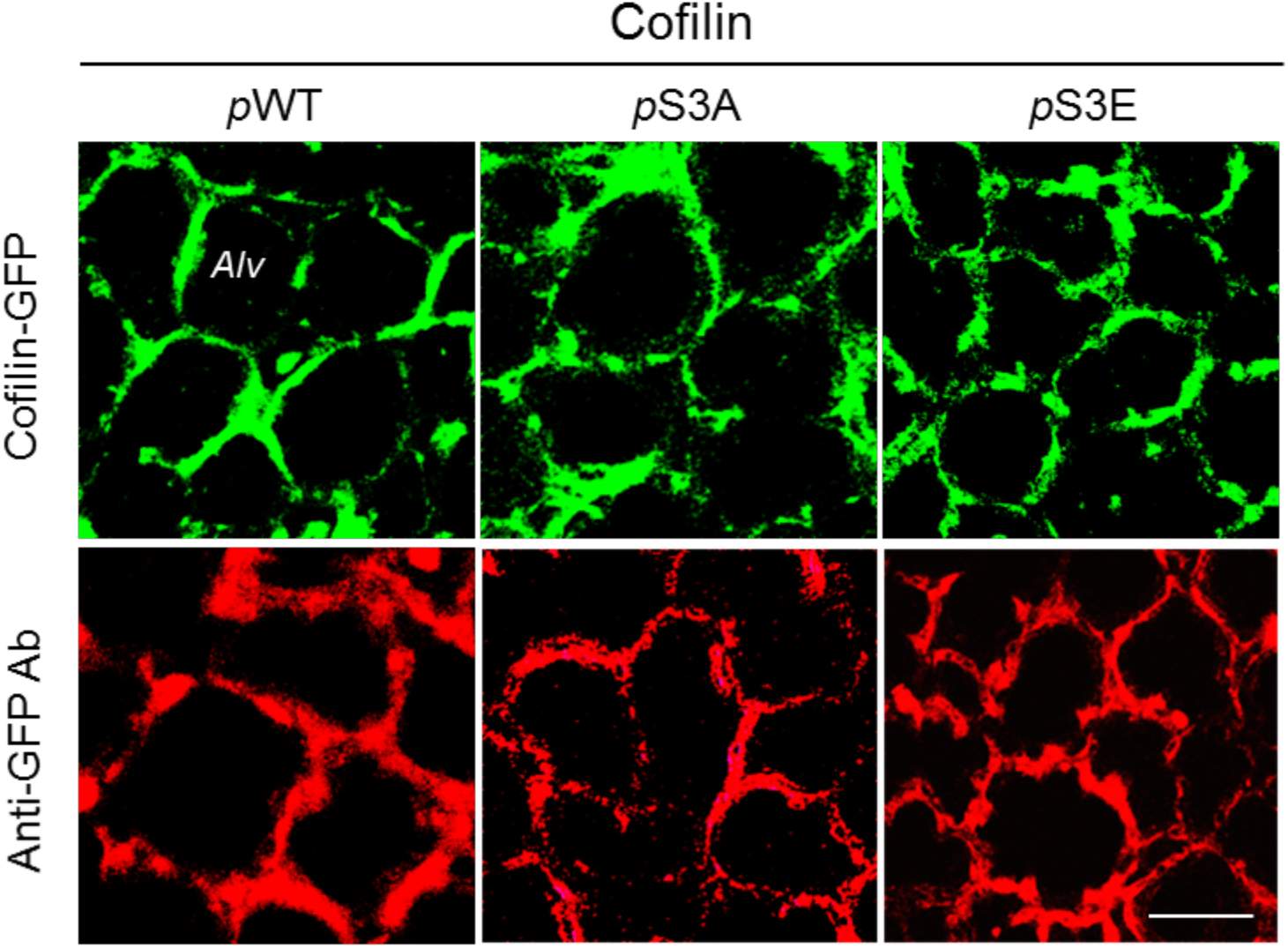
Expression of cofilin mutants in alveoli. Confocal images show fluorescence of the expressed GFP-cofilin mutants (green) and anti-GFP immunofluorescence (red) confirming alveolar expression of the cofilin. Cofilin transfections were for wild-type plasmid (*p*WT), and for constitutively active (*p*S3A), or inactive (*p*S3E) cofilin mutants; *Alv*, alveolus. Scale bars, 50 μm. Replicated in 3 lungs.

**Supplemental Figure 4.**
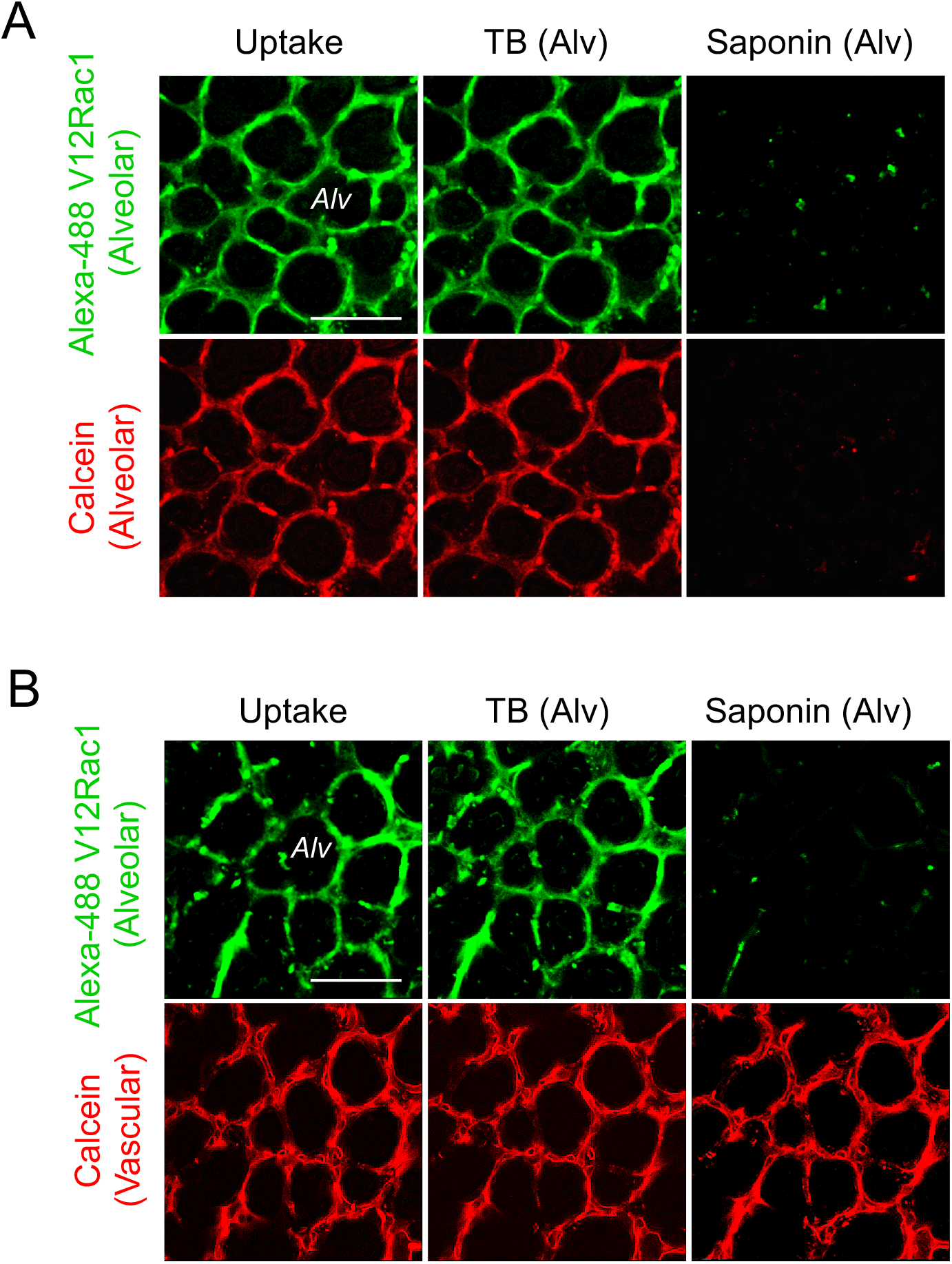
Rac1 uptake by alveolar epithelium. Confocal images show epithelial internalization of TAT-conjugated V12 Rac1 (upper panels, green) (**A, B**). Cytosolic dye, calcein (red) delineates the alveolar epithelium (**A**) and vascular endothelium (**B**). Alveolar infusion of trypan blue (TB) did not quench green fluorescence, confirming V12 Rac1 was internalized. Alveolar infusion of saponin decreased calcein fluorescence in alveoli (**A**), but not in microvessels (**B**), indicating V12 Rac1 was internalized by alveolar epithelium alone. *Alv*, alveolus. Scale bars, 50 μm. Replicated in 2 lungs.

**Supplemental Figure 5.**
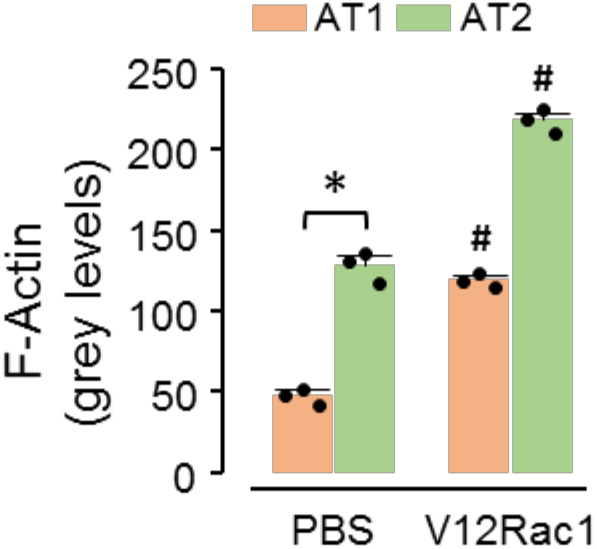
Effect of Rac1 on alveolar epithelial cells. Bars show fluorescence quantifications of AT1 and AT2 actin after the indicated treatments. Each bar is mean±SE for 8 quantifications per lung, in 3 lungs. \**p* < 0.05 as indicated, #*p* < 0.05 versus corresponding baseline using 2 tailed *t* test. The dots are means of determinations for each lung.

**Supplemental Figure 6.**
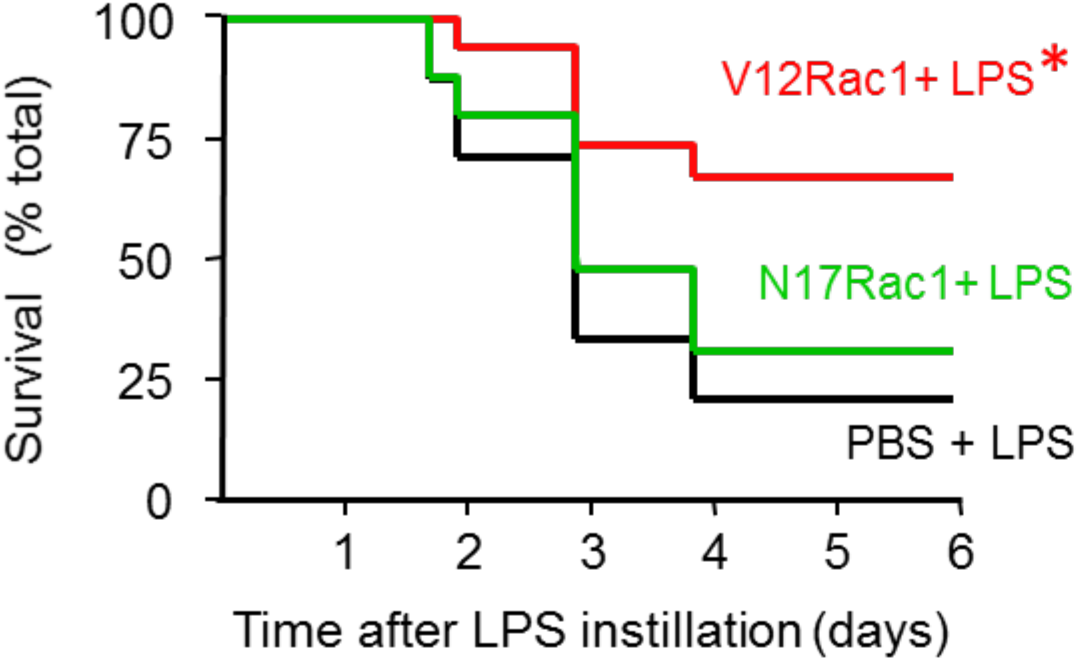
Mouse survival. Kaplan-Meier plots show survival in mice pre-treated i.n. with TAT-Rac proteins, followed 30 mins later with intranasal LPS (LD80) instillation. *n*=15 mice in each group, *p<0.05 versus LPS using long-rank test.

**Supplemental Figure 7.**
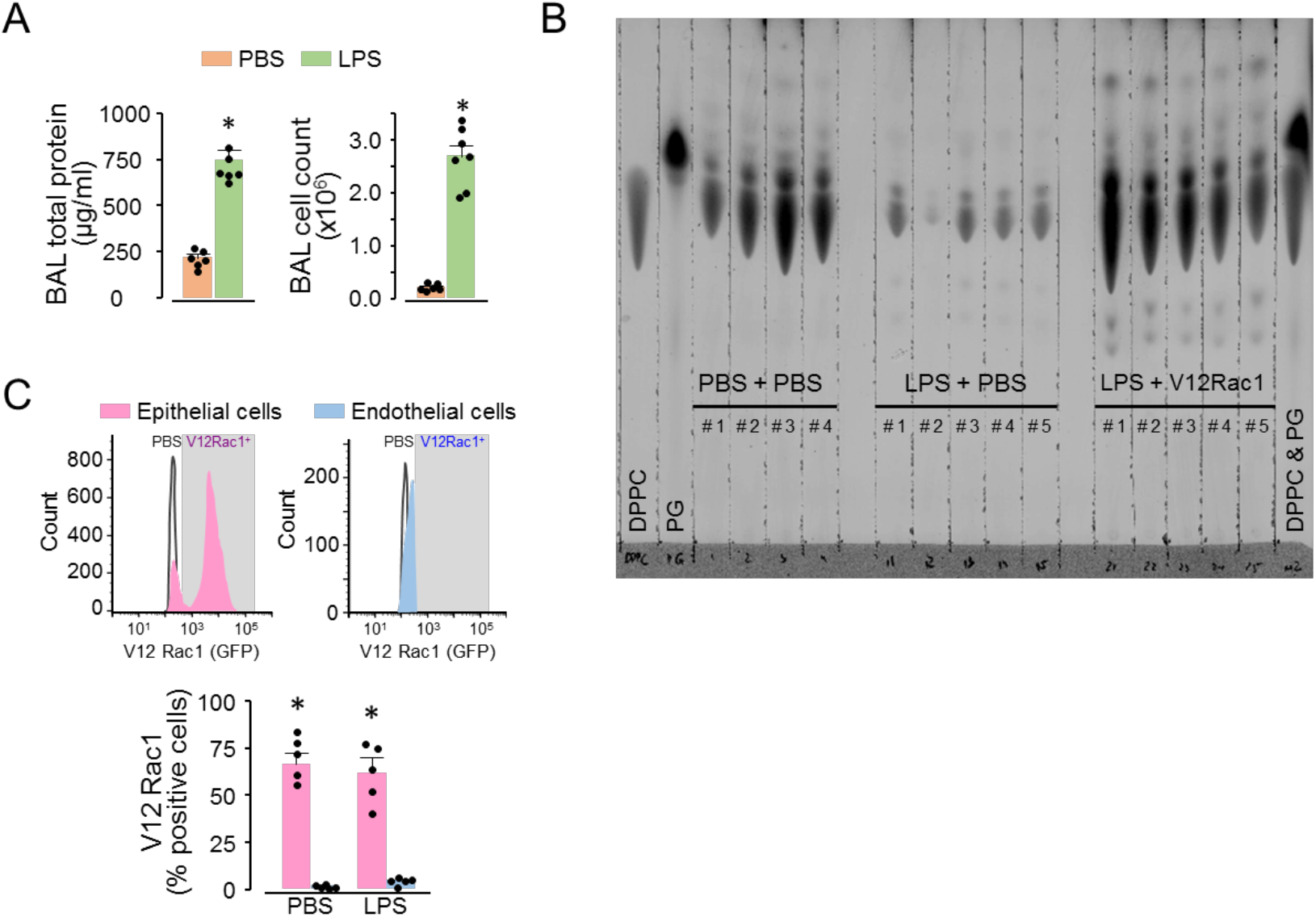
Global Rac1 uptake and Thin-Layer Chromatography (TLC) analysis of BAL. **(A-C)** Mice were given intranasal instillation of LPS at LD80 or PBS. **A** shows BAL total protein and leukocyte counts 4 h after LPS. Mean±SE, \**p*<0.05 vs PBS using 2-tailed *t* test. *n*=6 in PBS, and *n*=7 in LPS groups. Each dot shows data for a single lung. In **B** and **C**, intranasal TAT-V12Rac1 was given 4h after the LPS or PBS instillations, then after 72 (**B**) or 1 (**C**) hour, lungs were removed for BAL phospholipid and flow cytometry analyses, respectively. **B** shows an image of a TLC plate for phospholipid analyses. *n=*5, except PBS (*n*=4). Histograms in **C** show fluorescent V12Rac1 uptake (grey box). Bars are percentages of V12Rac1-positive cells in the indicated cell types. Mean±SE, *n*=5 lungs for each group, \**p*<0.05 using ANOVA with Bonferroni correction. Each dot shows data for a single lung.

**Supplemental Figure 8.**
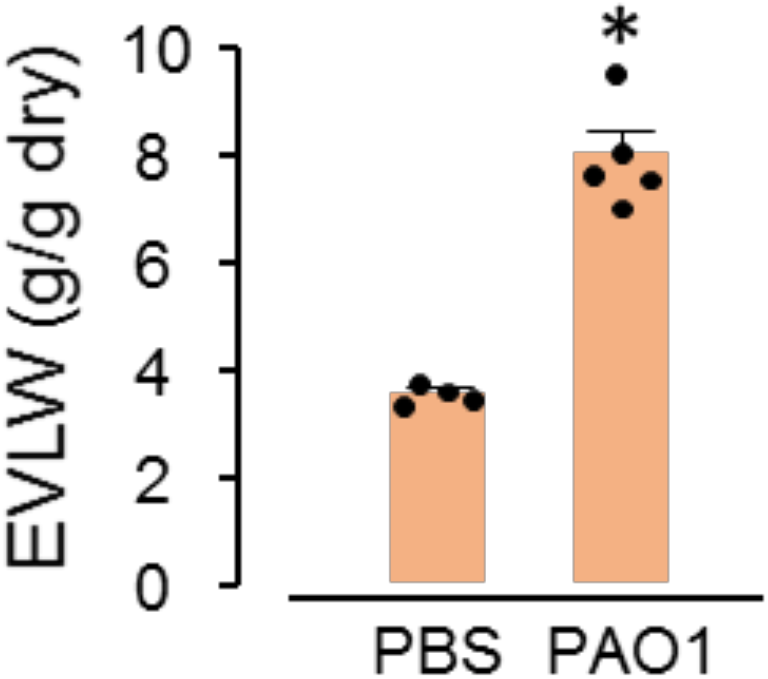
Quantification of pulmonary edema. Bars show blood-free extravascular lung water (EVLW) determined 4 h after *i.n.* instillation of *P. aeruginosa* (PAO1) or PBS. Mean±SE, *n*=4 for PBS and *n*=5 for PAO1. \**p*<0.05 *vs*. PBS using 2-tailed *t* test. Each dot shows data for a single lung.

